# When complex movement yields simple dispersal: behavioural heterogeneity, spatial spread and parasitism in groups of micro-wasps

**DOI:** 10.1101/2022.09.14.507936

**Authors:** Victor Burte, Melina Cointe, Guy Perez, Ludovic Mailleret, Vincent Calcagno

## Abstract

1. Understanding how behavioural dynamics, inter-individual variability and individual interactions scale-up to shape the spatial spread and dispersal of animal populations is a major challenge in ecology. For biocontrol agents, such as the microscopic *Trichogramma* parasitic wasps, an understanding of movement strategies is also critical to predict pest-suppression performance in the field.
2. We experimentally studied the spatial propagation of groups of parasitoids and their patterns of parasitism. We investigated whether population spread is density-dependent, how it is affected by the presence of hosts, and whether the spatial distribution of parasitism (dispersal kernel) can be predicted from the observed spread of individuals.
3. Using a novel experimental device and high-throughput imaging techniques, we continuously tracked the spatial spread of groups of parasitoids over large temporal and spatial scales (eight hours; and six metres, ca. 12,000 body lengths). We could thus study how population density, the presence of hosts and their spatial distribution impacted the rate of population spread, the spatial distribution of individuals during population expansion, the overall rate of parasitism and the dispersal kernel (position of parasitism events).
4. Higher population density accelerated population spread, but only transiently: the rate of spread reverted to low values after four hours, in a “tortoise-hare” effect. Interestingly, the presence of hosts suppressed this transiency and permitted a sustained high rate of population spread. Importantly, we found that population spread did not obey classical diffusion, but involved dynamical switches between resident and explorer movement modes. Population distribution was therefore not Gaussian, though surprisingly the distribution of parasitism (dispersal kernel) was.
5. Even homogenous asexual groups of animals were shown to develop behavioral heterogeneties over a few hours. Explorer individuals were responsible for most parasitism and dispersal, and determined spatial spread and density-dependent dispersal. We showed that simple Gaussian dispersal did not emerge from simple diffusion, but rather from the interplay of several non-linearities at individual level. This suggests expectations from classical diffusion theory may not hold generally to active dispersers. These results highlight the need to take into account behaviour and inter-individual heterogeneity to understand population spread in animals.

## INTRODUCTION

The movement and dispersal of animals through their environment is a fundamental process that affects a range of biological scales, from individual fitness, to population dynamics and spatial spread, and, ultimately, gene flow and evolution (Bowler & Benton, 2005; Clobert et al., 2012; Ronce, 2007). Movement is a complex phenomenon arising from individual behaviour and decision-making, but is also shaped by population processes and landscape characteristics. As a result, the study of animal movement is intrinsically a multiscale question, requiring the integration of methods and concepts from several disciplines in behaviour, ecology and physics (Lewis et al., 2013; Nathan et al., 2008; Turchin, 1998). Continued developments in the technologies available to track animals and analyse their movement patterns provide unprecedented amounts and quality of data to dissect the mechanisms governing animal movement at different scales (Branson et al., 2009; Nathan et al., 2022). In particular, studies of movement in the lab have benefited from progress in image acquisition and computer vision, opening new avenues in the tracking of entire populations and automatic detection of life-history events and behaviours (Dell et al., 2014; Robie et al., 2017). Long restricted to single individuals and small temporal and spatial scales, experimental studies now extend to entire groups and to larger and larger scales (Fronhofer et al., 2015; Kuefler et al., 2012).

Understanding the causes of variability in dispersal and their consequences for ecological dynamics is and has long been, a major goal of ecological research. For instance, understanding how one can upscale detailed processes at the level of individuals into patterns at the population scale is a prime ecological question (Sutherland et al., 2013; Spiegel et al., 2017). Bridging studies of individual behaviour with population-level studies of dispersal is in this regard a major challenge, and a crucial one if we are to better predict the response of populations to changing environments and landscape characteristics (Holyoak et al., 2008). From an applied perspective, understanding the spatial spread of groups of animals has high significance. For instance, in biological control agents used in inundative releases for crop protection, the capacity to disperse from the release points and thoroughly search the environment is recognized as a key phenotype in determining field efficiency, and also controls the optimal spatial disposition of releases in a field (Chapman et al., 2009; Cônsoli et al., 2010).

The experimental study of animal movement is however challenging, because of the spatial and temporal scales involved, and because of the great variety of movements and behaviours individuals can have. At a small scale, detailed observations of individuals can help understand their strategic decision making and how movement decisions are influenced by local environmental or social factors (Cronin, 2009; Calcagno et al., 2014; Kreuzinger-Janik et al., 2022). At large scales, the study of population spread and dispersal patterns inform us on the outcome of all individual decisions and on average quantities crucial for population dynamics and evolution, such as rates of spread or levels of gene flow (Dahirel, Bertin, Haond, et al., 2021; Morales et al., 2010; Petrovskii & Morozov, 2009). However, systems are still scarce for which the two types of data are simultaneously available and can be integrated, explicitly linking behavioural and population processes (Michelangeli et al., 2022). This scarcity has both practical and technological reasons, and it is, ultimately, one goal of movement ecology to fill this gap (Nathan et al., 2022).

In this study, we propose a novel experimental approach to study population spread in a small insect, and to dissect the effects of population density and resource distribution on movement, dispersal and total resource consumption. Our study system is a minute parasitoid wasp from genus *Trichogramma*, one of the smallest insect species known and a major biological agent (Cônsoli et al., 2010). Taking advantage of its minute size (<0.5 mm), of a dedicated experimental system and of high-definition camera sensors, we were able to track in continuous time the propagation of dozens of individuals at ecologically relevant temporal and spatial scales: six metres (ca. 12,000 body lengths) and eight hours. This spatial scale is comparable to the typical dispersal distance covered in the field by these insects in one day, as they mostly explore by walking and are incapable of directed flight (Gardner & Hoffmann, 2020). As for the temporal scale, most movement and parasitism take place within the first day in this short lived species. Furthermore, *Trichogramma* are egg parasitoids, so that their hosts are immobile and their distribution can be experimentally controlled with great precision. The location of parasitism events is also directly connected to dispersal in the strict population genetics sense (i.e. place of offspring births) as the entire development occurs within hosts. Finally, our study-species is thelytokous, so that mature females emerge for parasitized hosts and immediately start their search for hosts to parasitize. This dramatically simplifies the diversity of life-stages and types of movements that occur in natural populations, making this more amenable to being reproduced in a laboratory setting. This system thus offers unique characteristics for the experimental study of spatial propagation and dispersal.

We used our experimental system to introduce groups of individuals and study the effects of population density, of the presence or absence of hosts, and of their spatial distribution (diffuse versus clumped) on the dynamics of population spread and parasitism. All these factors are known to potentially have a strong impact. Increasing the density of individuals would often increase dispersal, as was observed in several species (Rosenberg et al., 1997; Kuefler et al., 2012; Kreuzinger-Janik et al., 2022) including a related *Trichogramma* wasp (Mills & Lacan, 2004). Some organisms need to reach a density threshold to disperse in fragmented habitats (Morel-Journel et al., 2016). Such a positive density-dependence of dispersal might occur at high population densities, whereas the opposite (negative density-dependence dispersal) can also be observed at low densities (Fronhofer et al., 2015). The presence of resources, in our case hosts, can also strongly impact the rate of population spread, in one direction or the other. On the one hand, the presence of resources can elicit more locally-restricted search strategies, reduce the incentive to disperse into another part of the habitat, and divert activity time away from movement, factors all contributing to decrease the rate of spread (Kreuzinger-Janik et al., 2022). On the other hand, the presence of resources could also stimulate activity and promote movement, which might increase the rate of spread. Both population density and the presence of resources control the frequency of inter-individual interactions, as the latter can act as aggregation points. Such a direct effect is supplemented by the indirect effect of population density on the rate of resource depletion; the two factors might thus interact in their effect on population spread. For instance, time allocation and oviposition strategies were both directly and indirectly affected by the presence of conspecifics in parasitic wasps (Mohamad et al., 2015; Robert et al., 2016). These few examples demonstrate how intricate the connections between movement, parasitism and dispersal can be.

In this article, we will precisely quantify the rates of population spread observed in different conditions, as measured by mean squared displacements and diffusion coefficients. We will test the assumptions of classical diffusion theory, in terms of rates of spread and of the spatial distribution of individuals. We will also quantify the dispersal kernel (spatial distribution of parasitism events), and confront it to the observed dynamics of population spread. We will show that even in such a simple and controlled experimental setting, population spread does not conform to classical diffusion theory. Our homogeneous groups of clonal individuals develop a dynamical behavioural heterogeneity that affects the rates of spread and the distribution of individuals. We show that population spread can be very well described by a model of heterogeneous diffusion with dynamical switches between “explorer” and “resident” movement modes. We show that, surprisingly, this relatively complex movement dynamics results in a simple, Gaussian, dispersal kernel. We argue that the simple dispersal kernel occurs not because of a simple diffusive underlying movement, but rather because of several non linearities in the dynamics of parasitism and resource depletion. Our results highlight that dispersal and population spread need not be related in a simple linear manner, and that seemingly diffusive dispersal kernels do not necessarily indicate simple diffusive spatial spread, or vice versa. This stresses the importance of taking into account behavioural heterogeneity and dynamics in order to predict the spatial spread and dispersal of groups of animals.

## MATERIALS & METHODS

### Study species and rearing conditions

Experiments were conducted with *Trichogramma cacoecieae*, sampled in 2014 near Mallemort (43.74387N, 5.12603E; Vaucluse, France). Species identity was confirmed by morphological analysis and COI barcoding. Live individuals from this strain are available (ID code PMBIO1) from the Biological Resource Center in our institute. *T. cacoeciae* is a strictly thelytokous species, whose females give birth to genetically identical daughters by clonal reproduction (Cônsoli et al., 2010). Insects were reared on UV-sterilised eggs of *Ephestia kuehniella* (the common laboratory host for trichograms, also used for sampling and experiments) with a 12:12 light:dark cycle, at temperatures switching between 25°C and 19°C, in order to get a development time of 14 days from host parasitism to adult emergence (Burte et al., 2022). All individuals used for experiments were 24 hour old and kept at 25°C in a glass tube with water and honey as a facultative food source prior to experiments.

### Experimental set-up and modalities

Individuals were released in the centre of a six metres (630 cm) long tunnel (1cm wide and 9mm high). At the scale of our insects (ca. 0.4 mm long), 630 cm represent about 12,000 body-lengths, and even the width and height of the tunnel exceed their visual perception range (reactive distance is about 4 mm; (Bruins et al., 1994)). To be able to cover the entire tunnel with digital cameras, the tunnel was folded into a double spiral, thereby fitting inside a rectangular area of 60 by 40 cm (Fig. 1). The double-spiral arenas were carved into styrofoam (Depron) plates 9mm thick, placed on a sheet of translucent paper on top of a glass plate, and sealed with another glass plate above. The device was uniformly lit from below with a light box (xxx lumens, colour temperature 5400K). All experiments took place in a dark room with controlled temperature (24°C ±1°C) and humidity (60% ±10%). To control for potential topological effects, half of the spirals were levogyrous (turning to the left, as in Fig.1) and half dextrogyrous (turning to the right). Further details on this experimental set-up and its performance can be found in a companion paper (Cointe et al. 2022).

**Figure 1:**
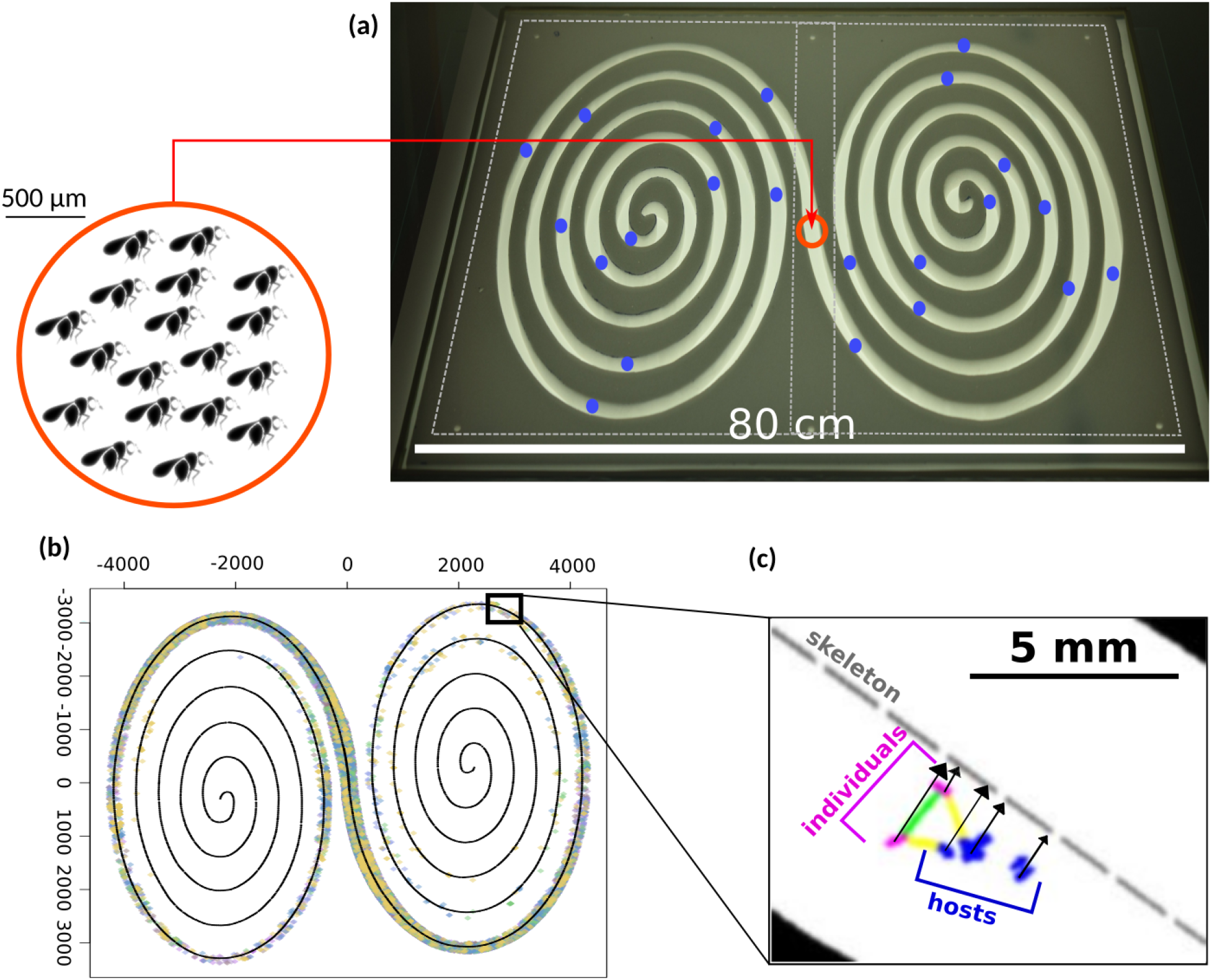
experimental set-up. Experimental set-up used for spatial propagation experiments. **(a)** A six-metre long tunnel was folded into a double spiral to fit in a 80×50 cm rectangle. The red dot figures the central point of release, and blue dots represent the location of host patches (Clumped modality: one patch every 30 cm). **(b)** Using high definition cameras image analysis techniques, the distribution of individual insects could be followed every minute for eight hours. **(c)** Individual insects and host eggs were located, and projected orthogonally on the tunnel skeleton to obtain their linear distance from the release point. Pairwise distances between individuals at any time, and distances to the closest host, were also computed.

At the beginning of an experiment, at 9 am, the desired number of individuals were briefly put at 9°C (to slow them down prior to transfer) and introduced at the centre of a double-spiral. Upon release they scattered over about 10 cm along the centre of the tunnel. The device was then immediately sealed with the top glass plate. Individuals were let propagate in the double-spiral tunnel for eight hours. At 5pm the experiment was stopped and individuals were manually removed. The device was then disassembled and cleaned with 70% ethanol before further use.

The number of individuals introduced was 25± 5 (Low Density treatments) or 70± 10 (High Density treatments). In addition, experiments were performed in which *E. kuehniella* host eggs were also put in the arena. In this case, host eggs were placed individually on the bottom sheet of paper, with a fine brush and a little water. Two types of host distributions were created: Diffuse (one egg every 5cm along the spiral tunnel, the first eggs 2.5 cm on each side of the centre), or Clumped (one patch of 6 eggs every 30cm along the spiral, the first 15 cm each side of the centre). Eggs were approximately centred between the two walls of the tunnel. All experiments with hosts were conducted at High Density (70 ± 10 individuals). The two host distributions thus had on average the same host density (120 eggs in total per arena). In those treatments the parasitism status of each egg at the end of the experiment was recorded: the bottom paper sheet was stored for five days at 25°C, after which parasitized hosts turned black and could be counted (see (Burte et al., 2022; Morel-Journel et al., 2016)).

Overall we conducted four types of experiments, randomised in time, between September 2018 and March 2019: (i) Low Density (20 replicates), (ii) High Density (20 replicates); (iii) High Density + Diffuse hosts (20 replicates); and (iv) High Density + Clumped eggs (22 replicates). On a given day, up to two experiments were run simultaneously in the same room.

### Image analyses and population tracking

After the introduction of individuals, high-definition pictures of the entire double-spiral arena were taken every minute throughout the experiment, i.e. for eight hours. In order to successfully visualise individual micro wasps (0.5 mm long) at this large spatial scale, two high-definition (full-frame 36 Mpx sensors) digital cameras were used synchronously, each centred on one end of the spiral, and covering half of the latter. This definition of 72 Mpx per minute ensured that *Trichogramma* appeared as dark objects of about 8 pixels on the light background. Images were stored in uncompressed RAW format, representing (2×480 images), i.e. about 220GB of image data per replicate.

Images were subsequently (off-line) analysed with a custom pipeline written in ImageJ (Schneider et al., 2012). Details and source code are available in Cointe et al. (2022). Raw images were converted to Tiff RGB, imported to ImageJ, and transformed to grayscale using a linear combination of brightness and saturation channels that maximised insect detection. Analysis then consisted in computing for each image a background image (as a moving window of 20 minutes), subtracting the background, thresholding, and particle analysis, yielding an ensemble of particles as putative trichogram detections. In parallel, the geometry of the double-spiral was resolved from the first image. In particular, the skeleton of the tunnel was reconstructed. Raw particle data and data on spiral geometry were then exported to R (R Core Team, 2020). Based on coordinates, size, shape and position relative to tunnel boundaries, a filter was applied to remove spurious detections. Data from the two cameras were realigned and combined to obtain one set of particles along the entire spiral. Finally, each particle was projected orthogonally on the skeleton of the double-spiral to be attributed a (signed) linear distance from the centre along the tunnel, and pixel distances were converted to centimetres. Additionally, if hosts were used, the exact location of each host egg was recorded by manual pointing on the first image in ImageJ, and the data was exported to R. Details of the analysis pipeline, as well as the source code, are available in Cointe et al. (2022).

Several tests were conducted to validate this pipeline (Supporting Information; see also Cointe et al. 2022). Among them, we checked that the number of detections accurately predicted the number of individuals introduced. With the settings retained, we could identify every minute on average 33% of all individuals introduced. This detection rate has several causes including false negatives (detections incorrectly discarded as artefacts) and non-detections (some individuals could not be seen on the walls where they occasionally wandered). For a subset of replicates, we also manually spotted all individuals that could be seen on the native images, and confronted their number and position with those detected with our pipeline. We could confirm that the overall detection rate was about 40%, consisting in a visibility rate of about 80% (i.e. 20% of individuals were not visible in any particular moment even by visual inspection) and in a methodological detection rate of about 50% (less-than-perfect algorithm). Most importantly, we checked that the detection rate was constant along the entire length of the tunnel, and that the spatial distribution of individuals and propagation metrics were accurately quantified by the method (Supporting Information; see Cointe et al. 2022). Finally, in none of our treatments did insects reach one end of the double spiral within eight hours, so that the groups can be conceived as dispersing freely in an open-ended tunnel, without boundary effects.

## Statistical analyses

For each replicate at every minute, we computed the Mean Squared Displacement (MSD), representing the total quantity of movement. In addition, we computed at every minute the population quantiles, i.e. the distance from the release point within which a certain proportion of individuals is found. Using the position of each host egg and spiral topology, we could also estimate for every insect at every minute its distance to the nearest wall, its distance to the nearest host egg, and the total number of individuals found on a given host egg at a particular time (see Fig. 1x). Furthermore, from parasitism data, the dispersal distance was calculated as the standard deviation of the distances from the release point to parasitized hosts. By regressing the average MSD on time, we estimated diffusion coefficients D (Turchin, 1998). 95% confidence intervals were obtained for the MSD, diffusion coefficients (D) and population quantiles using the bootstrap percentile method, with B=2000 resamplings. Comparisons between modalities for MSD values and population quantiles at given times were also performed using pairwise signed-rank tests with FDR corrections. Linearity of the increase of MSD over time was assessed by performing piecewise linear regressions and bootstrapping, using the R package segmented. The spatial distributions of individuals were compared to Gaussian through a comparison of percentiles (QQ-plot method). In addition, several candidate models were adjusted to the empirical distributions: the simple Gaussian model, Gaussian mixture models with two or three components, and finally a Student distribution representing a mixture of many Gaussian distributions with heterogeneous variance; Clark, 2007), all centred (i.e. with zero mean). These models were fitted by maximum likelihood and compared with AIC (Claeskens & Hjort, 2008). All analyses were performed in R (R Core Team, 2020). From the best models, parameters of interest were estimated, and the entire fitting procedure was repeated on B=600 bootstrap samples in order to estimate parameter uncertainty. All R code, with further details as comments, is provided as Supporting Information.

## RESULTS

### Population spread and the tortoise-hare effect

Population tracking provided, after filtering, a total of 761,409 insect detections, i.e. on average 162 detections per individual introduced in an experiment (raw data available in S.I.). In all treatments but the Low Density treatment, there was a relatively short (ca. 25 min) initial phase with slower propagation (Fig. 2), that we call a latency phase (see also Cointe et al. 2022).

**Figure 2:**
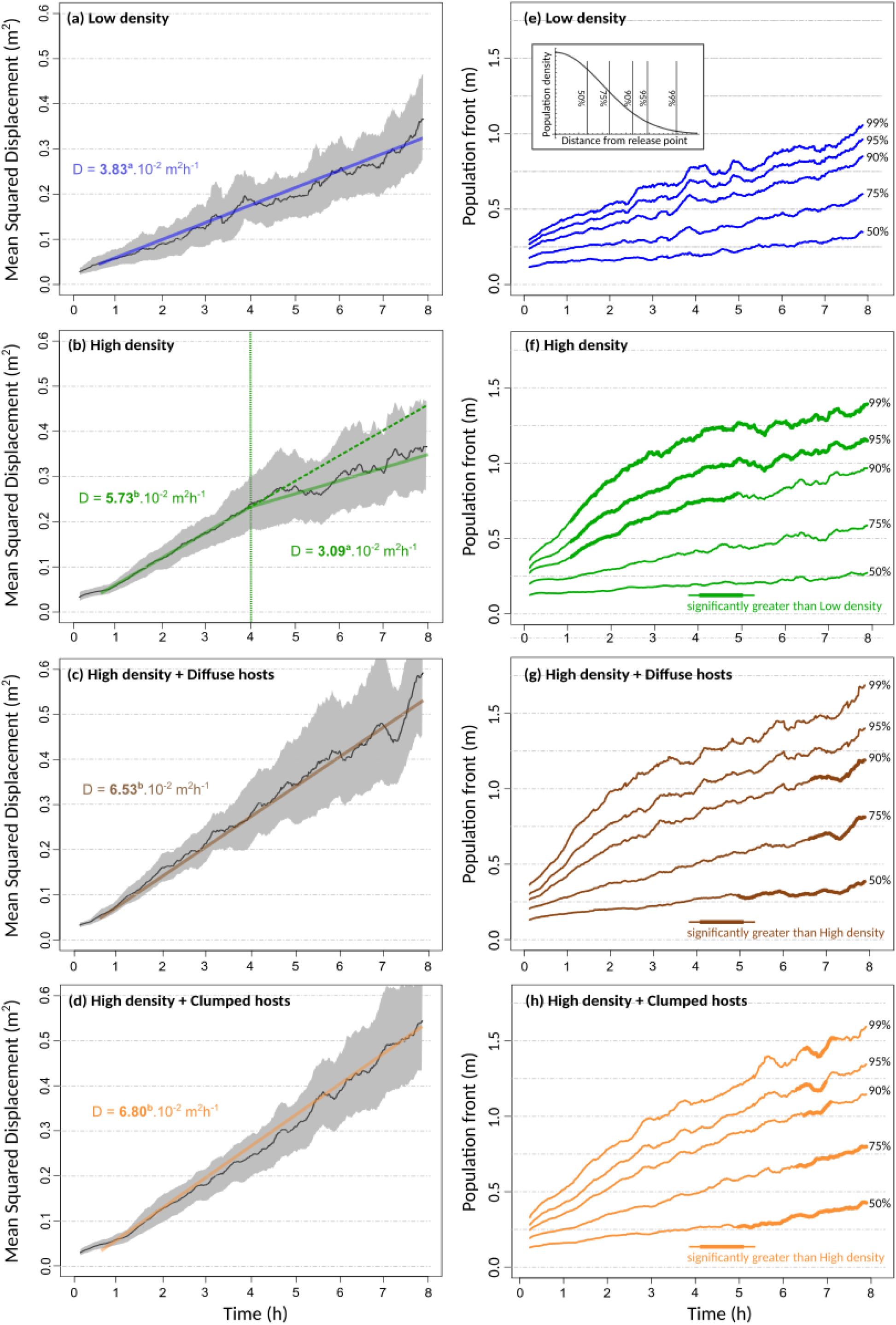
MSD and population fronts through time. Rate of population spread. **(a-d)** MSD as a function of time in the four experimental treatments. The solid curves show the average over replicates, with a moving window of 15 minutes. The grey envelope represents the corresponding 95% bootstrap confidence intervals. Linear regression lines are shown and their slopes (ie. diffusion coefficients D) are given. Significantly different values are indicated with different superscript letters. In the High Density treatment, a piecewise linear regression was significantly better supported, and thus two portions with two different diffusion coefficients are represented. The first 30 minutes were excluded from all regressions to avoid the effect of initial conditions and the latency phase. **(e-h)** The location (distance from centre) of population quantiles as a function of time. Bold sections represent significant pairwise differences, as indicated in the legend.

Overall, omitting the brief latency phase, the mean squared displacement (MSD) increased in a remarkably linear fashion with time (Fig. 2), in accordance with standard random walk and diffusion theory (Lewis et al., 2013; Turchin, 1998). A noticeable exception, however, was the High Density treatment, for which there was a marked slowing down of population spread after about four hours (Fig. 2B). In this treatment, the variation of MSD with time was significantly better fitted with a piecewise linear model, with a change in slope occuring at t=246 min. High Density treatments reached a coefficient of diffusion of D=5.73 m^2^h^-1^, which is much greater than the value observed in Low Density treatments (D=3.83 m^2^h^-1^). This indicates a strong positive density-dependence of the diffusion rate. However, this speed-up effect was only transient, and after about four hours the value of D in High Density treatment reverted to a low 3.09 m^2^h^-1^, not significantly different from the one in Low Density treatments (Fig. 2 a,b).

The transiency of density-dependence was not observed in the presence of hosts, for any spatial distribution (Diffuse or Clumped; Fig. 2). Rather, High Density treatments with hosts sustained elevated diffusion rates, not significantly different from the initial rate of spread observed in the absence of hosts, throughout the eight hours of the experiments (Fig. 2). As a consequence, the final MSD was much higher in the presence of hosts, reaching more than 0.5 m^2^ (Fig. 2c,d).

Spatial spread can be described more precisely through the direct measurement of population quantiles, i.e. the distance from the release point containing a given percentage of individuals. Figure 2 e-h represents the advancement of the core population (50% and 75% quantiles) and population front (90, 95 and 99% quantiles). At Low Density the tip of populations reached 1 metre at the end of the experiment, while High Density treatments went 50% farther, reaching 1.5 metre on average (Fig. 2). This is fairly consistent with the observed increase in diffusion coefficient. However, differences between treatments were not homogenous in different parts of the population. At High Density, population spread was faster than at low density, but the difference was driven only by the tips of the population (90-99% fronts), while no difference was observed in the core of the populations (50 and 75% quantiles; Fig. 2f). As a result, although at High Density the final total quantity of movement (MSD) was no greater than at Low Density (Fig. 2a,b), high density populations still covered longer distances than low density populations: the 90% and 95% population fronts remained significantly greater throughout the entire experiment (Fig. 2e-f). In effect, populations at High Density covered a distance similar to the treatment treatments with hosts (about 1.5m; Fig. 2f-h). Population fronts behaved similarly at High Density, with or without hosts, despite the strong difference in the total quantity of movement. This indicates that only the core of the populations caused the sustained rate of spread and higher MSD observed in the presence of hosts (Fig. 2g-h).

To summarise, a three-fold increase in initial population density caused a 1.5-fold increase in the rate of population spread (diffusion coefficient), at least transiently. However, in the absence of hosts, a marked deceleration in the second half of the experiments made the final MSD similar at High and Low Density. The initial boost provided by positive density-dependence thus did not pay off in the long run; we call this a “tortoise-hare” effect, by analogy with Aesop’s proverbial fable. These effects of population density on spatial propagation were mostly driven by population fronts (i.e. maximal distances reached), whilst, on the contrary, the effects of host presence manifested themselves mostly in the population core. Therefore, although the total quantity of movement at first sight obeyed classical diffusion, the population quantiles do not, and different processes operated in different parts of the populations.

### Distribution of individuals and evidence for heterogeneous diffusion

Consistent with results on population quantiles, the distribution of individuals differed markedly from the Gaussian distribution expected from classical diffusion theory (Fig. 3). Only at the very beginning of experiments did individuals adopt a Gaussian distribution, centred on the release point (Fig. 3). From about 30 minutes on, the distributions became strongly leptokurtic, with an excess of individuals at very short distances or very long distances (heavy tails), compared to Gaussian distributions with the same mean and variance. The observed leptokurticity could not be accounted for inter-replicate variation, since it persisted even after renormalizing each replicate to unit variance (S.I. Fig. S2). This was the case in all treatments and throughout the experiments, with distribution shapes changing dynamically.

**Figure 3:**
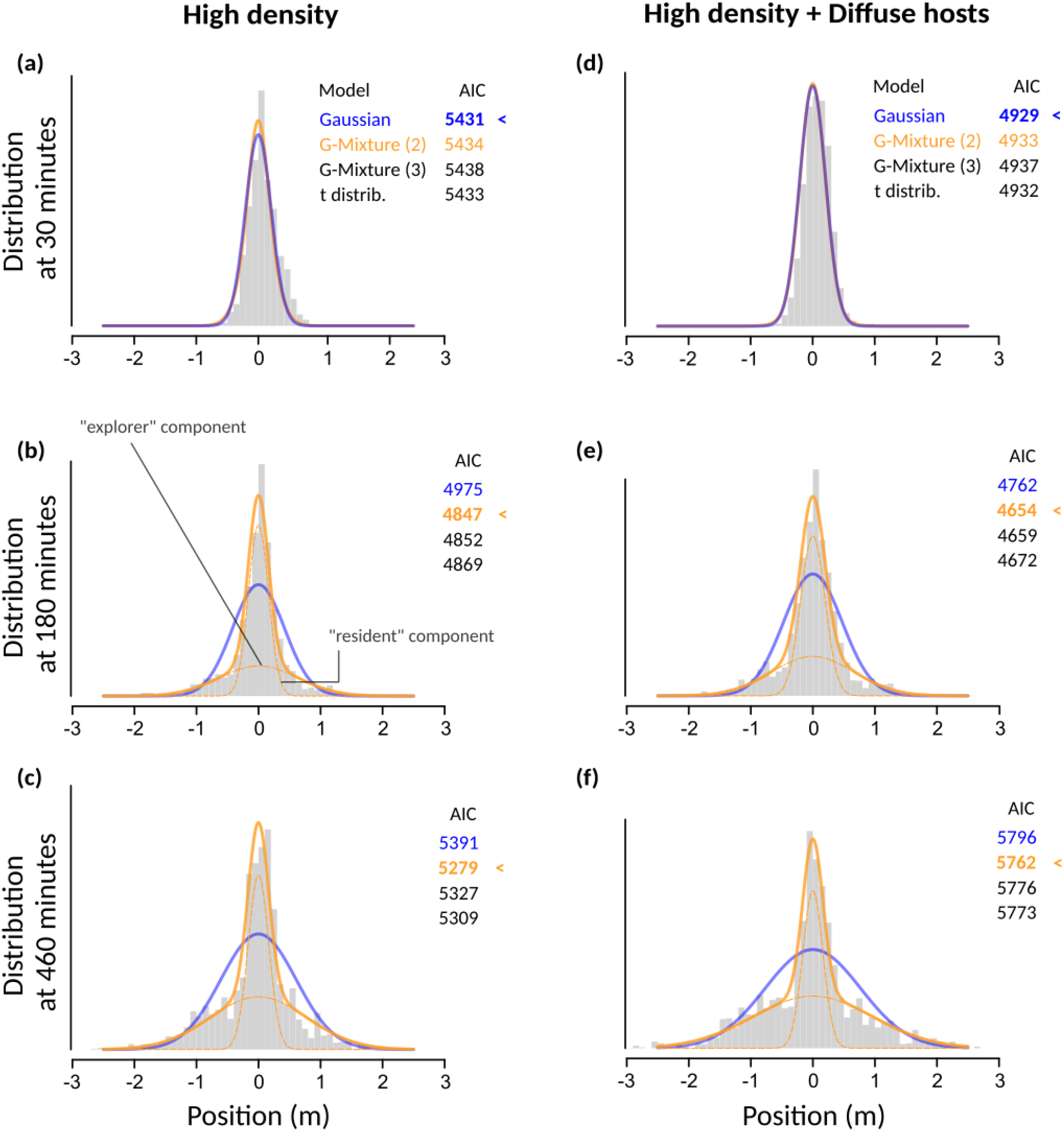
explorers and residents. The distribution of individuals during population spread. **(a-c)** High Density treatment and **(d-f)** High Density + Diffuse hosts treatments (patterns were similar in other treatments; see Figure S2 in S.I.). Histograms show the observed distribution of individual detections over all replicates, at three different times (15, 380 and 460 minutes). A time window of +/- 5 minutes, centred on the focal time, was used for better visualisation (but not for the statistics). Curves show the best fitting Gaussian (blue) and 2-component Gaussian mixture (G-mixture (2); orange) models. AIC values of Gaussian, G-mixture (2), G-mixture (3) and Student (t) distribution are also given in each case. Dashed lines represent the two components of the G-mixture (2) model, i.e. the narrow (“resident”) and broad (“explorer”) components.

One possible cause of leptokurtic distributions is individual heterogeneity, resulting in heterogeneous random-walks (Fraser et al., 2001; Lewis et al., 2013; Petrovskii et al., 2011). If each individual is diffusive, but there is variation in individual dispersal rates, the resulting population dispersal kernel, sometimes called a statistically structured kernel, would be fat-tailed (Petrovskii & Morozov, 2009). The fit of Gaussian mixture models revealed that a two-component mixture distribution fitted the observed data very well (Fig. 3). It statistically outperformed both the simple Gaussian distribution and alternative leptokurtic kernels (the 3-component Gaussian mixture and the t-distribution; Fig. 3). This was true for most times and treatments, except at the very beginning of experiments, where the distributions of individuals were indeed close to Gaussian or Student. This suggests that two types of individuals, differing in their movement strategies, coexisted in the populations. As one can see Fig. 3, the first component remained close to the central release point and spread very slowly, whereas the second component was much broader and diffused faster, constituting most of the population fronts. Such a bimodality in movement phenotypes has never been described in *Trichogramma*, especially at those large scales. By analogy with common terminology in the literature, we will call the first component the “resident” component, and the second the “explorer” component (Michelangeli et al., 2022).

From the method of mixture model fits, we can further infer the relative proportion of the two types, at every time, and compute the MSD for the two components individually (see S.I. for detailed Methods). Figure 4a-d shows the estimated proportion of resident and explorer individuals through time. In all treatments, proportions were not constant: populations initially comprised a majority of resident individuals, but the proportion of the latter steadily decreased in favour of explorer individuals. These dynamics differed across treatments. In the absence of hosts, the two types of individuals ended up being in roughly 50/50 proportion after eight hours (Fig. 4a-b). At High Density, the frequency of explorers increased faster initially, but then essentially plateaued, whereas the increase was more steady at Low Density. In the presence of hosts, the rate of conversion from resident to explorer strategies remained high for a longer period of time (Fig. 4e-f): the two types became equally frequent after only about three hours, and eventually explorers clearly outnumbered the other type, representing more than 60% of the population at the end of the experiments (Fig. 4c-d).

**Figure 4:**
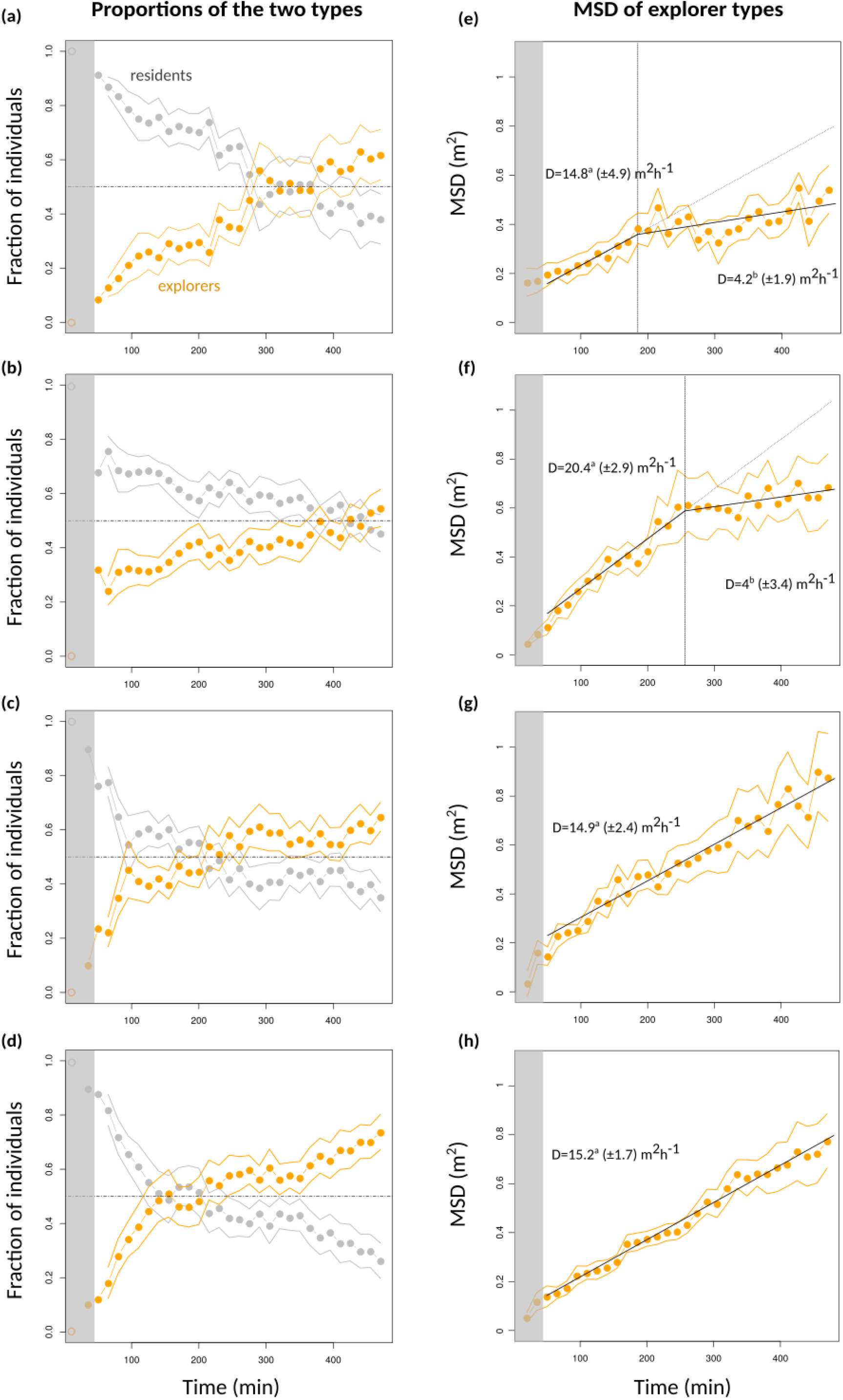
Proportion and MSD of explorers through time. Relative frequency and movement quantity of explorer individuals over the course of population spread. **(a-d)** Each dot represents the relative proportion of each component, as inferred from the G-mixture (2) model fit (see Fig. 3), every 15 minutes, over time windows of 10 minutes. Solid lines delineate the corresponding 95% bootstrap confidence intervals. Estimates for the first 30 minutes were unstable because they correspond to the latency phase (shaded part) in which the population distribution was still close to Gaussian (Fig. 3). The presumed initial states are shown as empty circles. **(e-h)** Each dot represents the inferred MSD for the explorer component, using the same methodology as in a-d. Best supported linear or piecewise linear regressions are represented, together with their slopes (diffusion coefficients) and associated standard errors. Significantly different values harbour different superscript letters.

In all treatments, explorer individuals initially adopted similarly high rates of spread of about 15 m^2^h^-1^ (Fig. 4e-h), but in the absence of hosts the diffusion coefficient of the explorer component eventually declined, reaching values of about 4 m^2^h^-1^, not significantly greater than resident strategists (see S.I.). The decline occurred earlier in low Density treatments than in High Density treatments, even though one must keep in mind that the estimation of these times is highly uncertain. One could thus argue that in the absence of hosts, explorer individuals that had initially dispersed far away effectively reverted to a resident strategy. In contrast, in the presence of hosts explorer individuals sustained the high diffusion coefficient throughout experiments (Fig. 4 g-h).

These behavioural dynamics explain the “tortoise-hare” effect and why the effects of population density were most expressed in population fronts (Fig. 2a,f): increasing density initially boosted the rate of conversion from resident to explorer individuals, and these by definition constituted most of the population fronts. However, the rate of conversion and the rate of spread of explorer individuals both stalled after a few four hours, whereas the changes were more gradual at low density. These dynamics similarly explain the effect of hosts on population spread: hosts hasted the rate of conversion to explorer individuals and, most importantly, made it persistent though time. This is why the effects of host presence were most visible in population cores (Fig. 2g-h): it is probably where switches from resident to explorer mostly took place, for it is where most resident individuals are located.

### Parasitism and the dispersal kernel

Consistent with results on spatial spread, Diffuse and Clumped host distribution treatments did not differ significantly in the fraction of hosts parasitized (p=0.62; 22% on average; Fig. 5a) or in the dispersal distance (p=0.38; 79.9 cm on average). The total parasitism rate culminated to an average 75% around the release point, and gradually declined over about two metres (Fig. 5a). Despite the strongly leptokurtic distribution of individuals during population spread, and thus the even more leptokurtic integrated population density, the distribution of parasitism was almost perfectly Gaussian (Fig. 5a). In that, it complied with the standard assumption of population biology (Turchin, 1998). The dispersal distance (standard deviation of the dispersal kernel) thus constitutes a dispersal coefficient in the usual sense (σ).

**Figure 5:**
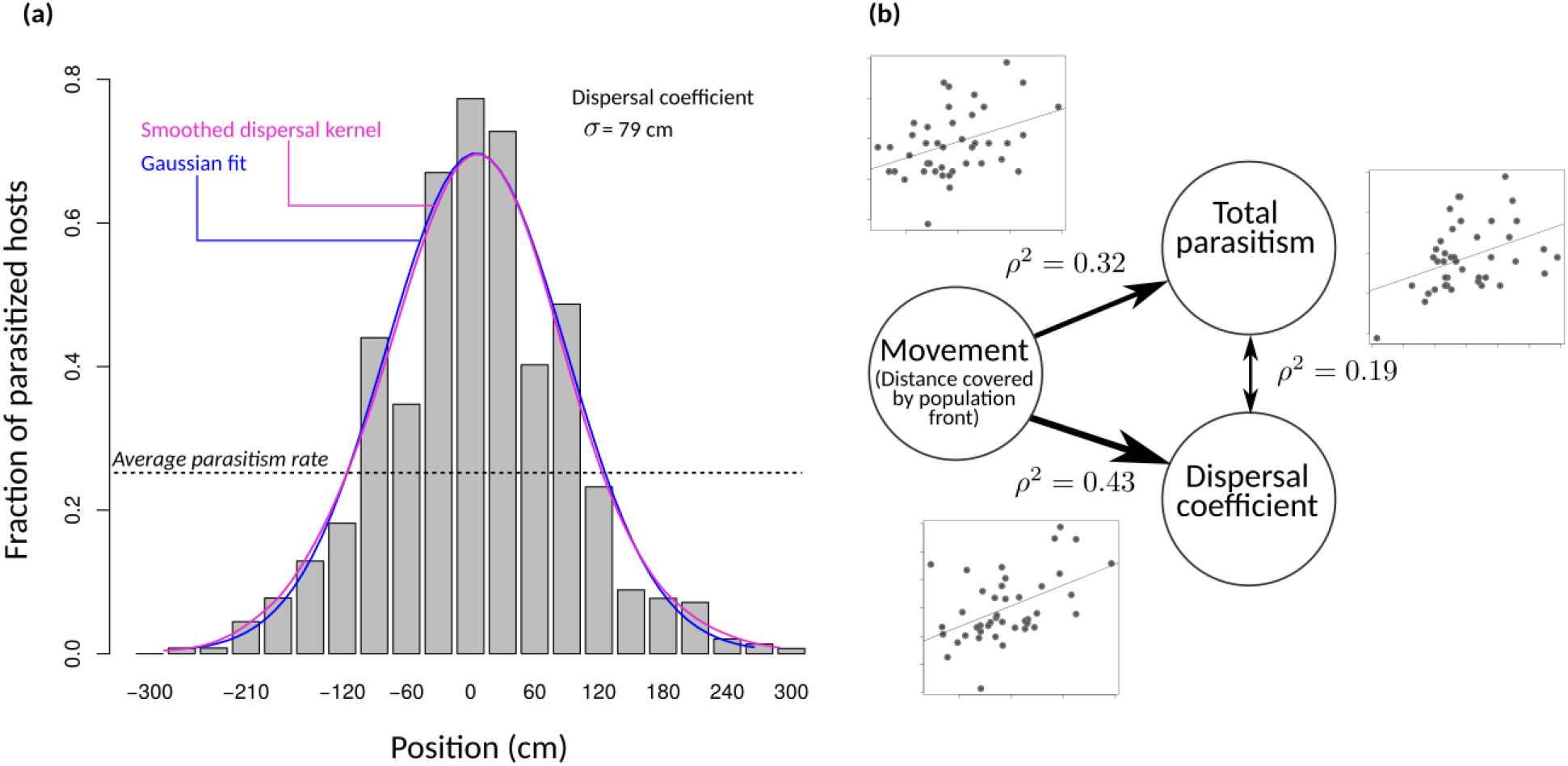
Dispersal kernel and parasitism. Parasitism and the dispersal kernel. **(a)** Dispersal kernel (fraction of hosts parasitized as a function of spatial position). The Diffuse and Clumped modalities having very similar distributions, the pooled distribution is shown, with bins of 30 cm (the distance between consecutive host patches in the Clumped modality). n=5,042 hosts. The continuous curves show the smoothed kernel (purple) and the fitted Gaussian distribution (purple). **(b)** Pairwise correlations between movement (distance covered by the 98% population front), total parasitism, and dispersal coefficient. Each point is one replicate (n=42 replicates). Spearman rank-correlations were used and the squared correlation coefficients are shown. Arrows have width proportional to the ρ^2^ values.

The spatial range over which significant parasitism occurred (3σ, or about 240 cm) greatly exceeded the 99% population front, which was located at about 150 cm (Fig. 2g-h). This suggests that population fronts made a very important contribution to parasitism. Confirming this, at replicate level, the position of the population front at the end of the experiment was a good predictor of the realised dispersal coefficient (r^2^=0.43; n=42; Fig. 5b). The best predictive performance was achieved using the 98% population quantiles, rather than other quantiles or the MSD (Fig. S3). The location of population fronts also predicted, to a lower extent, total parasitism (r^2^=0.32; Fig. 5b). Interestingly, the latter was rather weakly correlated to dispersal (r^2^=0.19; Fig. 5b).

### From movement to dispersal

How could individual distributions be strongly leptokurtic, yet produce an almost perfectly Gaussian dispersal kernel? Several non-linearities in the processes connecting individual movements to the location of parasitism events could explain this discrepancy. First, the fraction of individuals that were effectively on hosts (thus potentially parasitizing them) was not uniform, but was much higher at the population fronts at any time, reaching values as high as 60% (Fig. 6a). In the central parts of the population, this fraction quickly dropped to a baseline level of approximately 20%. Those spatial locations where most individuals were on hosts expanded as a wave as time passed (Fig. 6a). This indicates that individuals at population fronts presumably contributed much more to parasitism and dispersal. As a consequence, the huge density of individuals at the core of the populations did not translate linearly into parasitism pressure, for the decreased proportion of individuals that were effectively parasitizing hosts in these parts. This presumably reflects resource depletion and avoidance of already-parasitized hosts.

**Figure 6:**
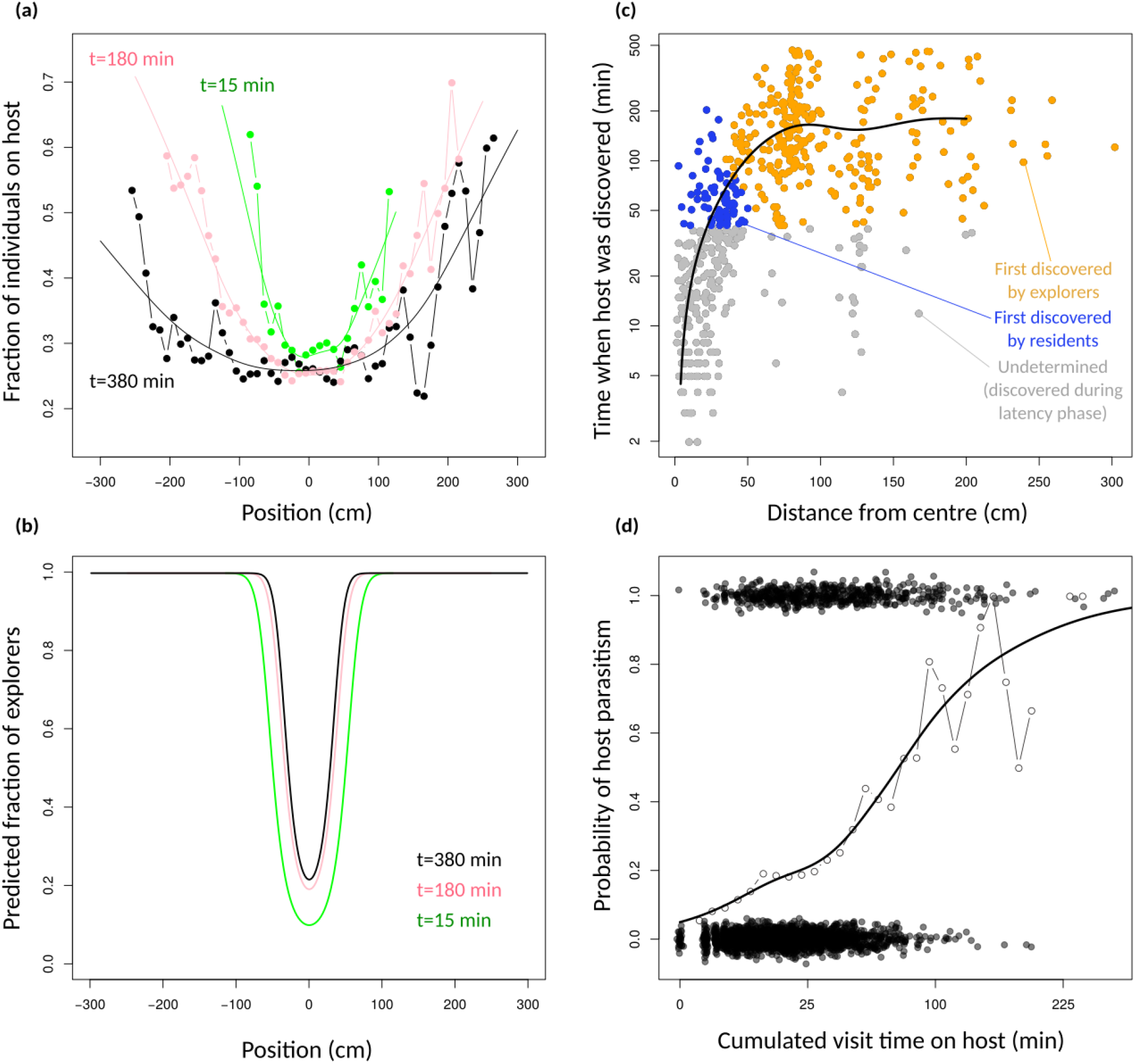
Connecting movement and parasitism. Connecting movement and dispersal. **(a)** The proportion of individuals that were in immediate proximity of a host, as a function of spatial position and at three different times: 15 min, 180 min and 380 min. Dots are sample proportions in bins of 10 cm. The continuous curve is the smoothed trend modelled as a general additive model. **(b)** The predicted proportion of individuals that were explorers at a given place and time. Predictions come from the mixture model fits shown in Figures 3 and 4. **(c)** The time at which hosts were first discovered, as a function of their spatial position (distance from centre). Each dot is a host. The continuous curve is the smoothed trend modelled with a general additive model. Dots were colored depending on whether they were most likely to have been discovered by an explorer individual (orange) or a resident individual (blue). Assignations were made using the predicted relative frequencies shown in panel b. Hosts shown in grey cannot be robustly assigned because they were discovered during the latency phase. **(d)** The probability that a host was parasitized as a function of the total visit time (cumulated time individuals have spent on it). Each dot is a host. The broken line represents the sample proportions, and the continuous curve is the smoothed trend modelled with a general additive model. In this figure, the Diffuse hosts treatment was used.

We can further confront the positions where most individuals were on hosts with the relative frequencies of explorer and resident individuals predicted from the Gaussian mixture fits (Fig. 6b). This reveals that at these locations, an overwhelming majority of individuals were explorers, except at the very beginning of the experiments. To substantiate this deduction, we can compute, for every parasitized host, the time at which it was first discovered, as a function of its position (FIg. 6c). From this and from Fig. 6b, we can then determine whether a particular parasitized host was most likely to have been discovered by a resident or an explorer individual. As can be seen in Fig. 6c, parasitized eggs beyond about 50 cm from the centre were most likely discovered by explorers, whereas hosts less than 50 cm away were most likely discovered by residents. This distance is to be compared with the dispersal coefficient (about 80 cm), and indicates that a majority (at least 56%) of all parasitism events can be attributed to explorer individuals. However, the contribution of the two types differed greatly between the centre of the dispersal kernel and its periphery.

Finally, while explorers at the population fronts reached most previously undiscovered hosts, they reached them at late times, and so did not have much time left to parasitize them. Host parasitism takes time in this species: processing time is usually about five minutes, and several visits may be needed before a host is accepted (Wajnberg et al., 2003). The probability of parasitism should therefore not increase linearly with the total visit time on a host: the so-called gain function is expected to be concave (Wajnberg et al., 2000; Calcagno et al., 2014). In line with these theoretical expectations, we found that the probability of host parasitism increased with total visit time, in a saturating fashion tending to one after about 200 min of cumulated visit time (Fig. 6d). This indicates that the density of individuals at the population fronts (the heavy tails of the individual distributions) also translated less-than-linearly into effective parasitism, for lack of sufficient host-processing time. Altogether, population density was most effectively converted into actual parasitism at intermediate positions, whereas this conversion was lower in central parts (because of resource depletion) and at the population tips (for lack of host processing time). We argue these two non-linearities, by decreasing the contribution of both central and extreme individuals, can explain the shift from a leptokurtic individual distribution to a simple Gaussian dispersal kernel.

## DISCUSSION

Bridging the gap between small-scale behavioural observations and large-scale population processes remains a prime challenge in ecology, and a requirement to alleviate disciplinary divides and improve our understanding and prediction capabilities. Methodological developments in the fields of sensors and computer-vision provide unprecedented power to take up this challenge, by allowing the acquisition of individual-level data in large groups and over large temporal and spatial scales (Holyoak et al., 2008; Nathan et al., 2022).

We used a novel experimental set-up and a dedicated imaging pipeline that allowed us to track the spatial spread of groups of insects along six metres and for eight hours. Six metres is considerable at the scale of our study system: it represents about 12,000 body lengths, or over 22 km if simply rescaling at human size. Even one centimetre, the width and height of our experimental maze, is still pretty big for *Trichogramma* insects: their reactive distance is about 4 mm (Bruins et al., 1994; Gardner & Hoffmann, 2020). For this reason, the inner volume of the double spiral set-up did not impose strong constraints on the movements of individuals: they could walk in 2D on all faces of the setting, which is their dominant form of exploration (Gardner & Hoffmann, 2020). They could also employ the occasional jumps and short undirected flights of the species (Cônsoli et al., 2010). Our hi-resolution imaging pipeline allowed us to locate every individual every minute, with a detection rate of about 40%, and thus to quantify not only the overall population spread, but also pairwise distances between individuals, distances to the closest host, the time spent parasitizing hosts, and many other individual metrics. We could thus address questions of population spread and dispersal with unique resolution.

We found that population spread was positively density-dependent: even though reared at similar densities, groups of insects released at higher density, as in the case in biocontrol programs, adopted a faster rate of spread (diffusion coefficient). This implies some quorum sensing at the stage of adults, which has never been demonstrated in this species, even though dispersal is known to depend on rearing density (Morel-Journel et al., 2016; Dahirel, Bertin, Calcagno, et al., 2021). The faster rate of spread took about 20-30 minutes to establish itself following introduction, after what we called a “latency phase”. This period is short on the timescale of our experiments, but quite long on the typical behavioural ecology timescale. It probably reflects an assessment period, during which individuals interact with one another and react to the local density (and, if any, to the presence of hosts). This interpretation is supported by the fact that in Low Density treatments with no hosts there was no detectable latency phase (Fig. 2a). It is also supported by the fact that the switching dynamics from sedentary to explorer movement mode occurred on a timescale of several hours (Fig. 4).

Interestingly, positive density-dependence was only transient in the absence of hosts. After about hours, the rate of spread reverted to a value no greater than the one observed at Low Density. The collapse of the rate of spread was actually so pronounced, that Low Density treatments, by sustaining a lower but more long-term diffusion coefficient, eventually caught-up with High Density treatments and achieved similar quantities of movement after eight hours. We called this a “Tortoise-Hare effect”, in reference to Aesop’s fable, according to which the slow and steady can win in the long run. This emphasises the difficulty to extrapolate short-term mechanisms on the longer term.

The faster initial rate of spread at High Density was caused by a faster rate of switching from resident to explorer mode, presumably for the higher rate of inter-individual interactions around the release point. However, by spreading faster, individuals quickly reduced the local density they experienced, as they diluted themselves across space and lowered the local levels of crowding. This in turn decreased the rate of switching, and eventually the individuals that had spread farther away reverted to a low diffusion coefficient, arguably switching-back to resident mode for lack of incentive. At Low Density, the rate of switching was initially lower, but as a consequence could be sustained for a longer period. All individuals were raised at the same density, and have similar experiences. The sole difference is their storage in tubes as groups of adults 24h before experiments: in principle, they may have perceived density differences at this stage, before the actual experiments. The initial frequency of explorers might thus have been higher in High Density treatments. Even though the trends all suggest the initial condition was zero explorers (Fig. 4), we cannot rule out this possibility. The existence of the latency phase, plus the fact that movement strategies changed dynamically over the course of experiments, also suggest that density-dependent effects occurred during the experiments, not before. In any case, the Tortoise-Hare effect shows that positive density-dependence is inherently self-defeating at large scale: the faster the initial spread, the stronger the decline in local population density and the suppression of the (local) fuel of density-dependence.

This was not true in the presence of hosts. The presence of hosts suppressed the Tortoise-Hare effect and allowed the elevated diffusion coefficient to be sustained in the long run. This was caused by a twofold effect. First, the rate of switching remained high for a longer period of time, and the overall proportion of explorer individuals reached higher values. Second, explorer individuals did not slow down even after several hours, i.e. presumably, did not switch back to a resident strategy. These two effects could be explained either by hosts acting as information-sharing points, at which individuals, even at low local density, can still perceive the presence of conspecifics, either directly by meeting on the hosts or indirectly through the perception of kairomones on the latter (van Alphen et al., 2003). Alternatively, the encounter of hosts could in itself act as a positive stimulus that enhances movement and spread, even though the visit of hosts and parasitism can have contrasted impacts on the short-term exploration strategy in parasitoids (Wajnberg et al., 2000; van Alphen et al., 2003). For instance, it has been reported that parasitoids could show decreased exploration activity when their egg-load decreases (Rosenheim & Rosen, 1991). In our case, egg load could be a source of individual heterogeneity in treatments with hosts, but the fact that explorers sustained high diffusion rates in those treatments, whereas they slowed down in the absence of hosts, rule this out as a possible explanation.

An important finding of this study is that the groups of individuals, even though uniform in terms of age, experience, and genetics, quickly developed into mixtures of two distinct movement strategies, that we called resident and explorer strategies. Inter-individual differences and personality traits are increasingly thought to be important determinants of population functioning (Dall et al., 2004; Dingemanse et al., 2010; Spiegel et al., 2017; Jolles et al., 2020). Variability in personality traits, boldness in particular, has been reported as a mechanism for leptokurtic distributions (Fraser et al., 2001). This consequence was confirmed in our study: the distribution of individuals at any time was not Gaussian, but strongly leptokurtic. Intra-population polymorphisms impacting movement and dispersal have been found in several organisms (Réale et al., 2007; Michelangeli et al., 2022), though not in *Trichogramma* so far. Intra-population variability in personality traits, specifically in activity-related traits, has been found in *Trichogramma*, but was of continuous nature (Lartigue et al., 2020). Our results showed that variation in movement was bimodal, not continuous, and also that it was dynamic in time, not static. Therefore it could not be attributed (solely) to cryptic genetic variation or inter-individual differences, but rather corresponded to behavioural plasticity, with alternative strategies that one individual can adopt depending on the events it experiences. Individual distributions could be fitted very well with a two-component mixture of Gaussian kernels, which offers quite interesting inference possibilities. This situation differs from most examples of variation reported so far. Obviously, we did not explicitly track individuals at a very small scale, so we cannot yet ascertain the exact behavioural determinants of the two strategies we indirectly detected. Further research will clarify these aspects, by supplementing our large-scale low-frequency tracking pipeline with small-scale video-tracking at different positions along the experimental tunnel.

While individuals in the resident mode had very low diffusion coefficient, it is important to stress that these individuals were still living and active individuals. Indeed, our imaging analysis pipeline disregards immobile or dead individuals. The estimated MSD of resident individuals, though very low and hard to estimate accurately, is clearly greater than the initial spread caused by the introduction of individuals in the arena. This confirms that some spread did occur even in resident mode.

The consequences of behavioural heterogeneity had much deeper implications than making individual distributions leptokurtic. Explorer individuals and the switching dynamics between alternative movement strategies was shown to be crucial in determining population distribution, dispersal, and total parasitism. First, the rate of switching from residents to explorers largely determined the effects of density-dependence, the Tortoise-Hare effect, and its suppression in the presence of hosts. Second, the diffusion coefficient of explorers determined the position of population fronts, which best explained the realised dispersal coefficient and total parasitism rate. Finally, explorer individuals discovered a majority of hosts, and were responsible for virtually all parasitism at distances beyond the dispersal coefficient. Yet despite all these complexities, the realised dispersal kernel proved to be almost perfectly Gaussian. This may seem surprising, especially considering that additional sources of heterogeneity often render dispersal kernels leptokurtic (Clark, 2007). We have suggested that different non-linearities in the processes linking movement to parasitism might provide the explanation. It therefore seems that the simplicity of the dispersal kernel, in our case, should not be taken as support for simple movement rules. It is likely that manipulating some conditions, such as the overall density and/or the abundance of hosts, could easily cause the apparent simplicity of dispersal to break apart. At this stage, this remains speculative, and further experiments and mechanistic mathematical modelling are needed to confirm our hypothesis.

In any case, our results suggest there can be quite a disconnection between parasitism and individual movement patterns. This has both fundamental and applied consequences. In biological control, the two are typically used interchangeably as proxies of dispersal, and it is common to extrapolate the 98% population front from the MSD, assuming Gaussian diffusion (Chapman et al., 2009). This method breaks apart with the sort of leptokurtic distributions we evidenced. More generally, predictions from classical diffusion theory should not be applied too broadly, even for quite simple active foragers such as *Trichogramma* wasps. Patterns of population spread and parasitism appear to be highly plastic, and therefore difficult to transfer across conditions. We suggest that improving our ability to understand this plasticity and our capacity to predict dispersal across contexts will come from better recognizing individual-level heterogeneity and the use of more behaviorally-mechanistic models.

## Acknowledgements

This research and VB’s PhD fellowship were funded by ANR (project TriPTIC, grant ANR-14-CE18-0002). MC acknowledges funding from project TrichoMove (INRAE/Université Côte d’Azur/Bioline Agrosciences).

## Authors’ contributions

VC, LM and VB designed the experiments. VB and GP performed the experiments. VB, MC and VC analysed the data. MC and VC wrote the manuscript with inputs from all authors.

## Data accessibility

Raw data tables and R scripts will be made available through Zenodo upon publication.

## References

Bowler, D. E., & Benton, T. G. (2005). Causes and consequences of animal dispersal strategies: Relating individual behaviour to spatial dynamics. Biological Reviews, 80(2), 205–225.

Branson, K., Robie, A. A., Bender, J., Perona, P., & Dickinson, M. H. (2009). High-throughput ethomics in large groups of Drosophila. Nature Methods, 6(6), 451–457.

Bruins, E., Wajnberg, E., & Pak, G. A. (1994). Genetic variability in the reactive distance in Trichogramma brassicae after automatic tracking of the walking path. Entomologia Experimentalis et Applicata, 72(3), 297–303.

Burte, V., Perez, G., Ayed, F., Groussier, G., Mailleret, L., van Oudenhove, L., & Calcagno, V. (2022). Up and to the light: Intra-and interspecific variability of photo-and geo-tactic oviposition preferences in genus Trichogramma. Peer Community Journal, 2, e3.

Calcagno, V., Grognard, F., Hamelin, F. M., Wajnberg, E., & Mailleret, L. (2014). The functional response predicts the effect of resource distribution on the optimal movement rate of consumers. Ecology Letters, 17(12), 1570–1579.

Chapman, A. V., Kuhar, T. P., Schultz, P. B., & Brewster, C. C. (2009). Dispersal of Trichogramma ostriniae (Hymenoptera: Trichogrammatidae) in Potato Fields. Environmental Entomology, 38(3), 677–685. https://doi.org/10.1603/022.038.0319

Claeskens, G., & Hjort, N. L. (2008). Model selection and model averaging. Cambridge Books.

Clark, J. S. (2007). Models for Ecological Data: An Introduction. Princeton University Press.

Clobert, J., Baguette, M., Benton, T. G., & Bullock, J. M. (2012). Dispersal Ecology and Evolution. OUP Oxford.

Cônsoli, F. L., Parra, J., & Zucchi, R. (2010). Egg Parasitoids in Agroecosystems with Emphasis on Trichogramma. https://doi.org/10.1007/978-1-4020-9110-0

Cronin, J. T. (2009). Habitat edges, within-patch dispersion of hosts, and parasitoid oviposition behavior. Ecology, 90(1), 196–207.

Dahirel, M., Bertin, A., Calcagno, V., Duraj, C., Fellous, S., Groussier, G., Lombaert, E., Mailleret, L., Marchand, A., & Vercken, E. (2021). Landscape connectivity alters the evolution of density-dependent dispersal during pushed range expansions (p. 2021.03.03.433752). bioRxiv. https://doi.org/10.1101/2021.03.03.433752

Dahirel, M., Bertin, A., Haond, M., Blin, A., Lombaert, E., Calcagno, V., Fellous, S., Mailleret, L., Malausa, T., & Vercken, E. (2021). Shifts from pulled to pushed range expansions caused by reduction of landscape connectivity. Oikos, 130(5), 708–724.

Dall, S. R. X., Houston, A. I., & McNamara, J. M. (2004). The behavioural ecology of personality: Consistent individual differences from an adaptive perspective. Ecology Letters, 7(8), 734–739. https://doi.org/10.1111/j.1461-0248.2004.00618.x

Dell, A. I., Bender, J. A., Branson, K., Couzin, I. D., de Polavieja, G. G., Noldus, L. P., Pérez-Escudero, A., Perona, P., Straw, A. D., & Wikelski, M. (2014). Automated image-based tracking and its application in ecology. Trends in Ecology & Evolution, 29(7), 417–428.

Dingemanse, N. J., Kazem, A. J. N., Réale, D., & Wright, J. (2010). Behavioural reaction norms: Animal personality meets individual plasticity. Trends in Ecology & Evolution, 25(2), 81–89. https://doi.org/10.1016/j.tree.2009.07.013

Fraser, D. F., Gilliam, J. F., Daley, M. J., Le, A. N., & Skalski, G. T. (2001). Explaining leptokurtic movement distributions: Intrapopulation variation in boldness and exploration. The American Naturalist, 158(2), 124–135.

Fronhofer, E. A., Klecka, J., Melián, C. J., & Altermatt, F. (2015). Condition-dependent movement and dispersal in experimental metacommunities. Ecology Letters, 18(9), 954–963.

Gardner, J., & Hoffmann, M. P. (2020). How important is vision in short-range host-finding by Trichogramma ostriniae used for augmentative biological control? Biocontrol Science and Technology, 30(6), 531–547.

Holyoak, M., Casagrandi, R., Nathan, R., Revilla, E., & Spiegel, O. (2008). Trends and missing parts in the study of movement ecology. Proceedings of the National Academy of Sciences, 105(49), 19060–19065.

Jolles, J. W., King, A. J., & Killen, S. S. (2020). The Role of Individual Heterogeneity in Collective Animal Behaviour. Trends in Ecology & Evolution, 35(3), 278–291. https://doi.org/10.1016/j.tree.2019.11.001

Kreuzinger-Janik, B., Gansfort, B., & Ptatscheck, C. (2022). Population density, bottom-up and top-down control as an interactive triplet to trigger dispersal. Scientific Reports, 12(1), 5578. https://doi.org/10.1038/s41598-022-09631-w

Kuefler, D., Avgar, T., & Fryxell, J. M. (2012). Rotifer population spread in relation to food, density and predation risk in an experimental system. Journal of Animal Ecology, 81(2), 323–329.

Lartigue, S., Yalaoui, M., Belliard, J., Caravel, C., Jeandroz, L., Groussier, G., Calcagno, V., Louâpre, P., Dechaume-Moncharmont, F.-X., Malausa, T., & Moreau, J. (2020). Consistent variations in personality traits and their potential for genetic improvement in the biocontrol agent Trichogramma evanescens. BioRxiv, 2020.08.21.257881.

Lewis, M. A., Maini, P. K., & Petrovskii, S. V. (2013). Dispersal, individual movement and spatial ecology. Lecture Notes in Mathematics (Mathematics Bioscience Series), 2071.

Michelangeli, M., Payne, E., Spiegel, O., Sinn, D. L., Leu, S. T., Gardner, M. G., & Sih, A. (2022). Personality, spatiotemporal ecological variation and resident/explorer movement syndromes in the sleepy lizard. Journal of Animal Ecology, 91(1), 210–223. https://doi.org/10.1111/1365-2656.13616

Mills, N. J., & Lacan, I. (2004). Ratio dependence in the functional response of insect parasitoids: Evidence from Trichogramma minutum foraging for eggs in small host patches. Ecological Entomology, 29(2), 208–216.

Mohamad, R., Wajnberg, E., Monge, J.-P., & Goubault, M. (2015). The effect of direct interspecific competition on patch exploitation strategies in parasitoid wasps. Oecologia, 177(1), 305–315.

Morales, J. M., Moorcroft, P. R., Matthiopoulos, J., Frair, J. L., Kie, J. G., Powell, R. A., Merrill, E. H., & Haydon, D. T. (2010). Building the bridge between animal movement and population dynamics. Philosophical Transactions of the Royal Society B: Biological Sciences, 365(1550), 2289–2301.

Morel-Journel, T., Girod, P., Mailleret, L., Auguste, A., Blin, A., & Vercken, E. (2016). The highs and lows of dispersal: How connectivity and initial population size jointly shape establishment dynamics in discrete landscapes. Oikos, 125(6), 769–777.

Nathan, R., Getz, W. M., Revilla, E., Holyoak, M., Kadmon, R., Saltz, D., & Smouse, P. E. (2008). A movement ecology paradigm for unifying organismal movement research. Proceedings of the National Academy of Sciences, 105(49), 19052–19059.

Nathan, R., Monk, C. T., Arlinghaus, R., Adam, T., Alós, J., Assaf, M., Baktoft, H., Beardsworth, C. E., Bertram, M. G., & Bijleveld, A. I. (2022). Big-data approaches lead to an increased understanding of the ecology of animal movement. Science, 375(6582), eabg1780.

Petrovskii, S., Mashanova, A., & Jansen, V. A. (2011). Variation in individual walking behavior creates the impression of a Lévy flight. Proceedings of the National Academy of Sciences, 108(21), 8704–8707.

Petrovskii, S., & Morozov, A. (2009). Dispersal in a statistically structured population: Fat tails revisited. The American Naturalist, 173(2), 278–289.

R Core Team. (2020). R: A Language and Environment for Statistical Computing. R Foundation for Statistical Computing. https://www.R-project.org/

Réale, D., Reader, S. M., Sol, D., McDougall, P. T., & Dingemanse, N. J. (2007). Integrating animal temperament within ecology and evolution. Biological Reviews, 82(2), 291–318. https://doi.org/10.1111/j.1469-185X.2007.00010.x

Robert, F.-A., Brodeur, J., & Boivin, G. (2016). Patch exploitation by non-aggressive parasitoids under intra-and interspecific competition. Entomologia Experimentalis et Applicata, 159(1), 92–101.

Robie, A. A., Seagraves, K. M., Egnor, S. E. R., & Branson, K. (2017). Machine vision methods for analyzing social interactions. Journal of Experimental Biology, 220(1), 25–34. https://doi.org/10.1242/jeb.142281

Ronce, O. (2007). How Does It Feel to Be Like a Rolling Stone? Ten Questions About Dispersal Evolution. Annual Review of Ecology, Evolution, and Systematics, 38(1), 231–253.

Rosenberg, R., Nilsson, H. C., Hollertz, K., & Hellman, B. (1997). Density-dependent migration in an Amphiura filiformis (Amphiuridae, Echinodermata) infaunal population. Marine Ecology Progress Series, 159, 121–131.

Rosenheim, J. A., & Rosen, D. (1991). Foraging and Oviposition Decisions in the Parasitoid Aphytis lingnanensis: Distinguishing the Influences of Egg Load and Experience. The Journal of Animal Ecology, 60(3), 873. https://doi.org/10.2307/5419

Schneider, C. A., Rasband, W. S., & Eliceiri, K. W. (2012). NIH Image to ImageJ: 25 years of image analysis. Nature Methods, 9(7), 671–675.

Spiegel, O., Leu, S. T., Bull, C. M., & Sih, A. (2017). What’s your move? Movement as a link between personality and spatial dynamics in animal populations. Ecology Letters, 20(1), 3–18. https://doi.org/10.1111/ele.12708

Sutherland, W. J., Freckleton, R. P., Godfray, H. C. J., Beissinger, S. R., Benton, T., Cameron, D. D., Carmel, Y., Coomes, D. A., Coulson, T., & Emmerson, M. C. (2013). Identification of 100 fundamental ecological questions. Journal of Ecology, 101(1), 58–67.

Turchin, P. (1998). Quantitative Analysis of Movement: Measuring and Modeling Population Redistribution in Animals and Plants. Sinauer.

van Alphen, J. J., Bernstein, C., & Driessen, G. (2003). Information acquisition and time allocation in insect parasitoids. Trends in Ecology & Evolution, 18(2), 81–87.

Wajnberg, E., Fauvergue, X., & Pons, O. (2000). Patch leaving decision rules and the Marginal Value Theorem: An experimental analysis and a simulation model. Behavioral Ecology, 11(6), 577–586.

Wajnberg, E., Gonsard, P.-A., Tabone, E., Curty, C., Lezcano, N., & Colazza, S. (2003). A comparative analysis of patch-leaving decision rules in a parasitoid family. Journal of Animal Ecology, 72(4), 618–626.

