## Supplementary Information for "When complex movement yields simple dispersal: behavioural heterogeneity, spatial spread and parasitism in groups of micro-wasps"

**Supporting Information for manuscript:**
### When complex movement yields simple dispersal: a high-throughput study of spatial spread and parasitism in parasitic micro-wasps

Burte, Victor[a1], Cointe, Mélina[a1], Perez, Guy[a], Mailleret, Ludovic[ab] & Calcagno, Vincent[a*]

[a] Université Côte d'Azur, INRAE, CNRS, Institut Sophia Agrobiotech, Sophia Antipolis, France
[b] Université Côte d'Azur, Inria, INRAE, CNRS, Sorbonne Université, Biocore, Sophia Antipolis, France

[1]equal contribution.

Content

1. More on the image analysis pipeline and detection performance p. 2
2. More on the leptokurticity of individual distributions p. 11
3. More on mixture-models fitting p. 13
4. MSD of the resident component p. 15
5. Predictive power of different population quantiles p. 16
6. Population distribution integrated over time p. 18

#### 1. More on the image analysis pipeline performance

The detailed experimental set-up and image analysis pipeline, together with in-depth validation of its performance, are available in the companion article Cointe et al. (2022). Even though the general elements presented in the latter apply, we provide below results that are specific to the actual experiments conducted in the present article.

##### A. Motivation

The results in the main document rely on the location of particles (assumed to be *Trichogramma* individuals) along the axis of the double spiral set-up. An important aspect of the results is the shape of the distribution of individuals along the spiral axis. First, it is found to be non-Gaussian (except at initial times), with a strong leptokurtic shape. Second, the MSD value (variance of the distribution at any time) is found to be linear through time, with the exception of "High Density" treatments : in this case, the increase of MSD through time decelerates considerably after about four hours of experiments.

Since the method is new, we cannot know for sure that the distribution of particles following image analysis really represents the actual distribution of *Trichogramma* individuals. One could argue that a non-Gaussian distribution, for instance, is an artefact: the method could distort a true Gaussian distribution into something leptokurtic. This is not particularly plausible, but we'd like to reject this possibility.

The only way to evaluate this is to obtain the "ground truth" for some images, and see how they compare to the distributions inferred from image analysis. This requires inspecting the images by eye and locating the Trichogramma individuals manually.

##### B. Methods

For seven replicates from the "High Density" treatment, we visually inspected the raw images, at three points in time : two hours, five hours, and seven hours. On each image, we manually tagged the *Trichogramma* individuals we saw using a colour dot in ImageJ. After this, the location of all tags on all images was automatically retrieved using an ImageJ macro. This corresponded to 7 replicates * 3 times * 2 sides (left or right half of the double spiral) = 42 images annotated, representing a total of 1,022 *Trichogramma* detections.

The manually annotated (x,y) coordinates of *Trichogramma* individuals were transformed into “distance from the centre of the spiral” by projecting them onto the skeleton of the spiral. This was done using the same data and R scripts as the main analyses. This provided the ground truth for this set of images, that could be compared to the corresponding results that we obtained using the automatic image analysis pipeline.
The main objective was to investigate the possibility that the patterns deduced from the automatic pipeline could be artefacts, and would not be found in the ground truth data. A secondary objective was to better quantify the performance of the automatic pipeline.

#### C. Results

##### C.1. Numbers of individuals

From the ground truth data, an average of 52 individuals was observed at any given time in a particular replicate. The average number of individuals introduced in these replicates was 66, suggesting we could see about 80% of individuals. The remaining 20% are presumably overlooked (even by eye, spotting *Trichogramma* is not easy at that scale) or were concealed behind wall sections (and thus absent from the image data). This proportion was the same at all times (2, 5 or 7 hours), indicating that we do not tend to “lose” individuals as time passes and distance from the introduction point increases..

##### C.2. Overall detection rate

From the image analysis pipeline, we recovered 529 detections, whereas the ground truth data yielded 1022 detections. This corresponds to an overall 52% detection rate, at the level of one particular minute in time (of course, the automatic pipeline returns one number every minute, and we can thus improve the effective detection rate by time averaging). The detection rate was variable across replicates and across times, ranging from 20% to 130%. This motivates the weighting of replicates by the number of detections (square root transformed) in our statistical analyses.

Note that if we combine the former 80% of individuals that are visible on the images, with the latter 50% detection rate, we obtain an overall detection rate of 40%. This is fully consistent with the estimate we obtained using only data from the image-analysis pipeline, across all treatments and replicates (40%; see main article).

##### C.3. Spatial distribution of individuals

In the ground truth data, the global distribution of individuals was markedly leptokurtic, as it is in the data from the automatic pipeline. Actually, for these seven replicates, the true distribution and the one inferred from image analysis are very similar (Figure 1).

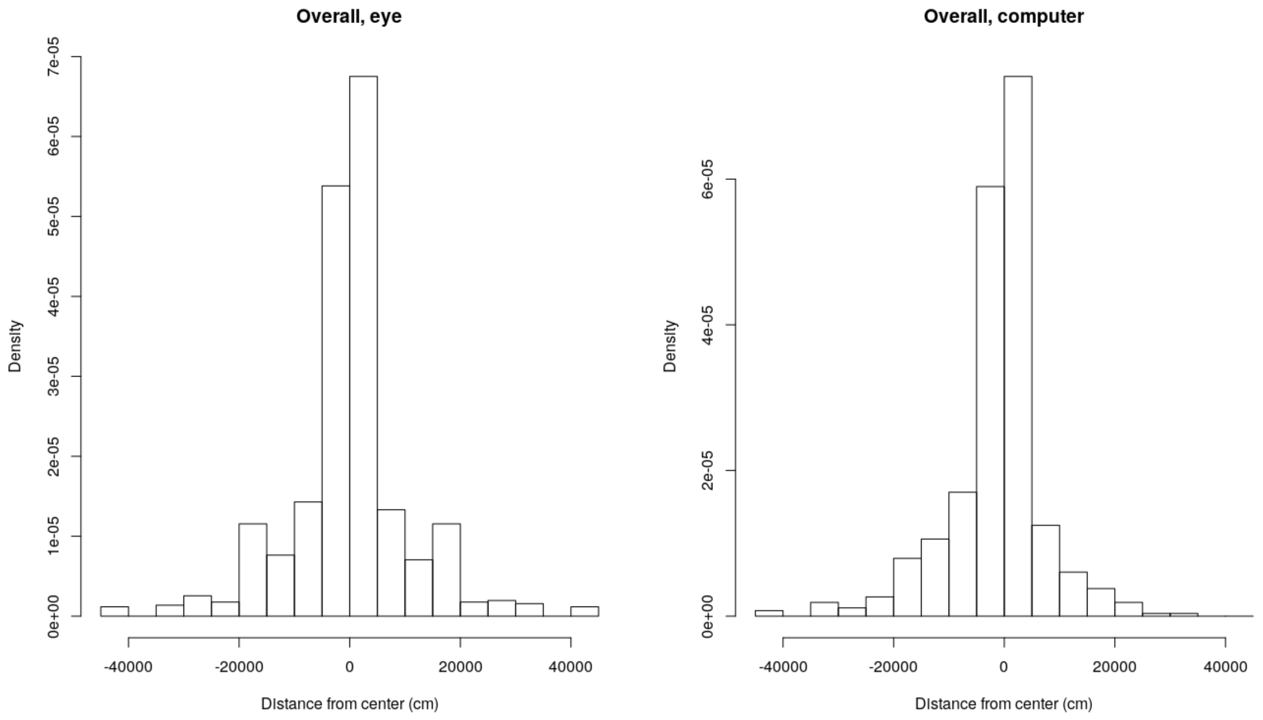


*Figure 1: Overall spatial distribution of individuals along the spiral axis (negative values refer to individuals inside the left branches, positive positions in the right branches). Left: from manual annotations (ground truth), right: from the automated image analysis pipeline (as used in the main article). This is for the 7 replicates that have been manually annotated, at times 120, 500 and 720 minutes. Note the similarity, and the fact that both are clearly leptokurtic.*

If we look in greater detail, at particular times, the same conclusion holds (Figure 2). After two hours, the two distributions are quite similar and both are clearly leptokurtic. Using Chi-square tests, the two distributions are actually not significantly different. The same at five and seven hours. Note that using the same test we can discriminate the distributions at different times, so lack of statistical power is not the only explanation.

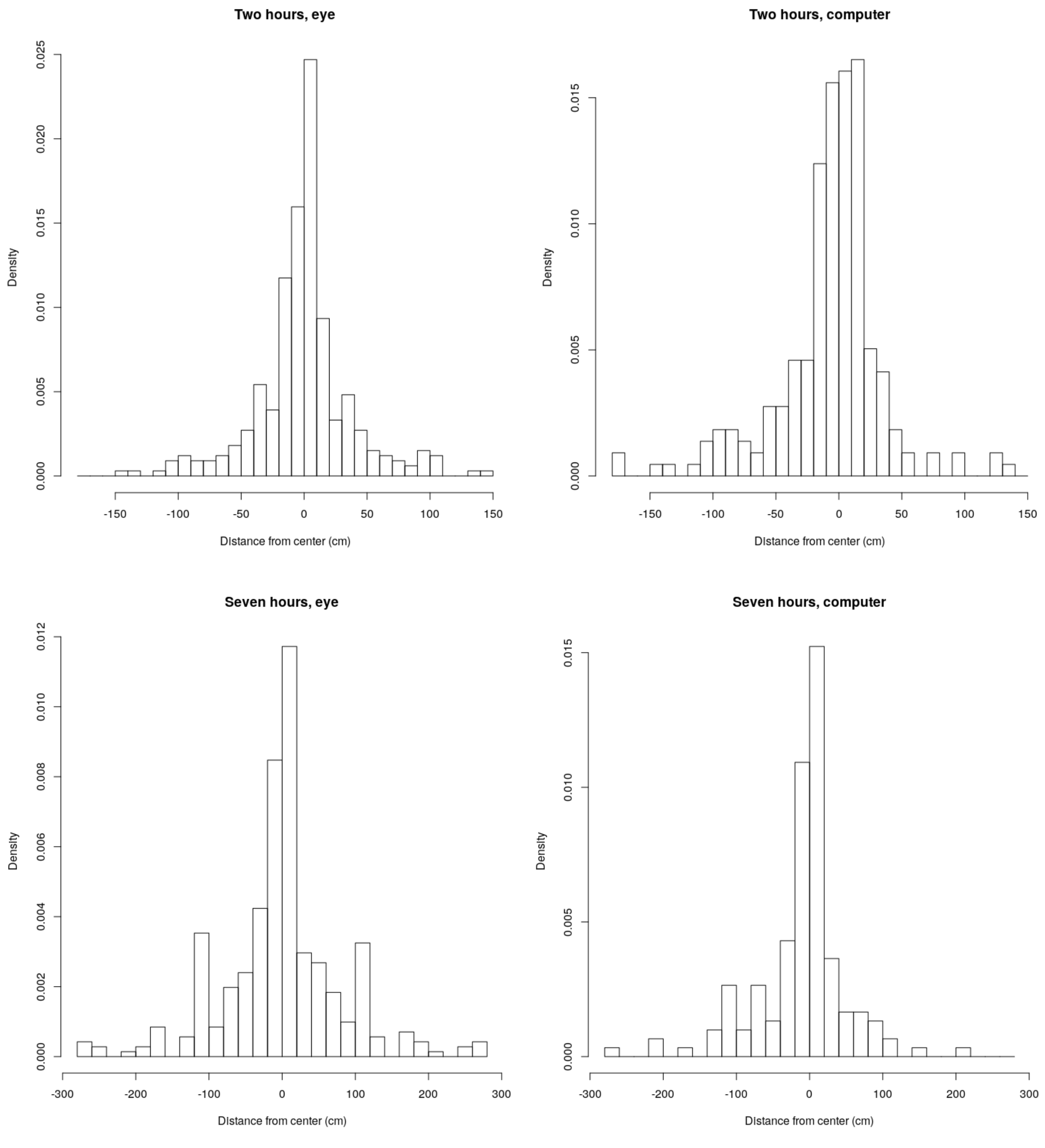


*Figure 2: Top: Spatial distributions at time 120 minutes. The two distributions are not significantly different from a Chi-squared test (p=0.26). Bottom: Spatial distributions at time 420 minutes. The two distributions are not significantly different from a Chi-squared test (p=0.06). Note the different x scales at the two times. At 300 minutes (not shown) the situation is similar and the two distributions do not differ either (p=0.09).*

If anything, we observe that the true distributions are even more leptokurtic than the ones from image analysis. This might be explained by a slight decrease in the detection rate as distance increases from the centre (Figure 3A). Note that these variations cannot create the leptokurtocity that we report. The observed distributions have a lack of density between 20 cm and one metre from the centre (see main article), and it is not where we observe a significant lack of detection power.
The impact of the decline in detection rate at far distances would be to underestimate the fraction of individuals that went very far, which would presumably reinforce our conclusions on leptokurticity (and matches the true distributions from the ground truth data).

Consistent with that, the detection rate slightly declines as time passes, as more and more individuals enter the remote locations where detection is less efficient (Figure 3B).

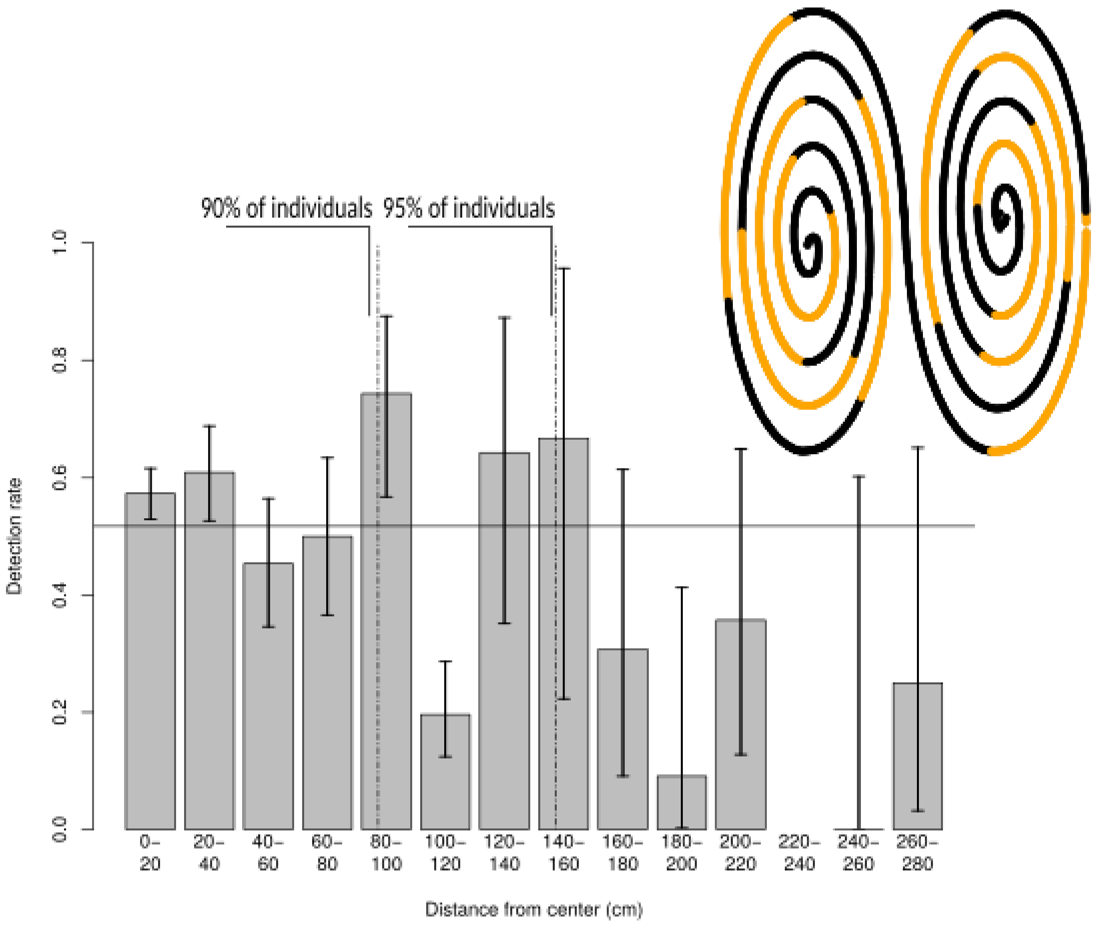
90% of individuals
95% of individuals
Detection rate
0.0
0.2
0.4
0.6
0.8
1.0
0– 20
20– 40
40– 60
60– 80
80– 100
100– 120
120– 140
140– 160
160– 180
180– 200
200– 220
220– 240
240– 260
260– 280
Distance from center (cm)

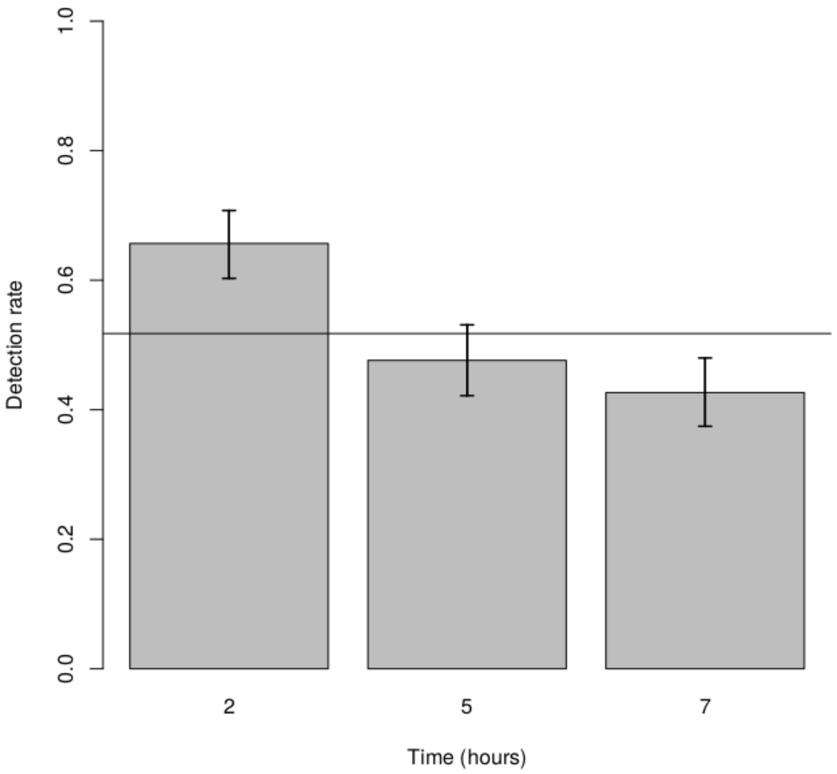
Detection rate
0.0
0.2
0.4
0.6
0.8
1.0
2
5
7
Time (hours)

*Figure 3: Top: Detection rate (number of detections from image-analysis pipeline divided by number in manual annotations) as a function of distance along the spiral. The spiral was divided in bins of 20 cm along each branch. The consecutive bins are shown in the insert drawing (consecutive bins are shown in alternating colours). Bottom: Detection rate as a function of time. The horizontal lines indicate the overall (irrespective of distance or time) detection rate.*

##### C.4 MSD values

The values of the MSD were almost perfectly correlated between the ground truth data and the data obtained from the automatic image analysis pipeline (Figure 4). In quantitative terms, the MSD tends to be underestimated at latest times, presumably because of the lower detection rate at farther positions, as explained in the previous section. However, the qualitative patterns, specifically the marked slowing down of MSD after four hours, as reported in the main article, are fully preserved, and they are well observed in the ground truth data.

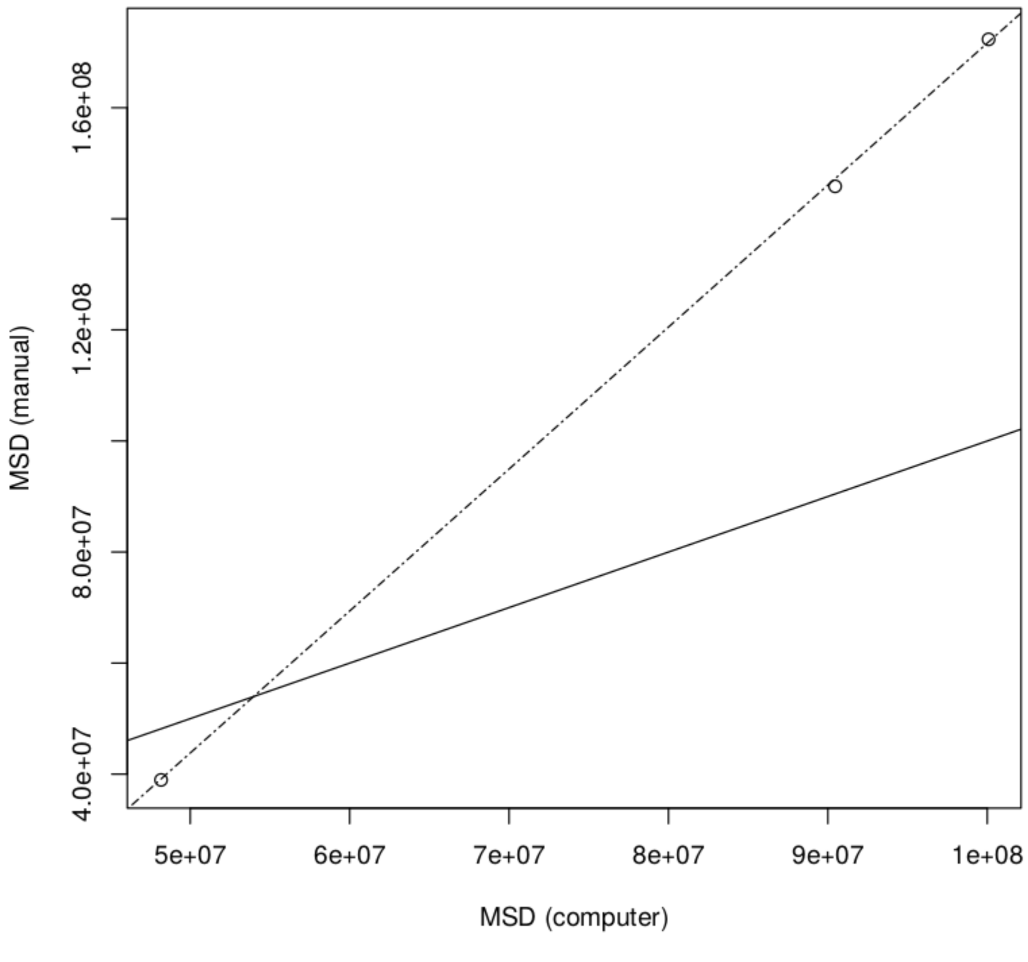

*Figure 4: MSD values at the three different times, based on the image-analysis pipeline (x-axis) or manual annotations (y-axis). Both MSD estimates increase through time (at 2, 5 and 7 hours, from left bottom to right top). The solid line is the 1:1 relationship. The dashed line is a linear regression (R2 = 0.99). Note the good agreement for MSD at the earlier time, and the slight underestimation of MSD from image-analysis, at the two latest times. Note also the perfect proportionality between the two estimates, and therefore the preservation of salient qualitative features. Specifically, the much slower increase in MSD between 5 and 7 hours, compared to that between 2 and 5 hours, i.e. the marked slowing down of spatial spread, is identical in the ground truth data.*

**D. Conclusions**

We conclude that the patterns reported in the main article are not brought-up by the image analysis pipeline, as they are also observed in “ground truth” data. The image analysis pipeline has good, though less-than-perfect, accuracy, and the detection rate is confirmed to be about 40% in any particular time (minute). The main characteristic of our pipeline is to underestimate the number of far distance movements, which may have some quantitative impacts: it may yield an underestimation of the “heavy-tailed-ness” of the spatial distributions, and it may thus cause some underestimation of the MSD values in the final parts of the experiments (i.e. for large MSD values; see Figure 4). These aspects have been improved in subsequent developments of the method (Cointe et al. 2022).

#### 2. More on the leptokurticity of individual distributions

The pooled distribution of individuals at a given time, across replicates, was found to be leptokurtic (Fig. 3 in main text). It is in principle possible that each replicate was in fact Gaussian, but variance heterogeneity across replicates generated a leptokurtic pooled distribution (in theory similar to a t-distribution). To investigate this hypothesis, we centred and renormalized (divided by the standard deviation) each replicate distribution individually, before pooling them. This procedure removes any effect of variance heterogeneity: if replicates were individually Gaussian, then the pooled distribution of renormalized positions would be Gaussian as well. Only if individual replicates are leptokurtic would the pooled distribution be leptokurtic. Leptokurticity can be best evaluated by using Quantile-Quantile plots; results are shown below (Fig. S1). They confirm the strong leptokurticity, even at replicate level. Considering that the conclusion is the same, and since natural distances are more interpretable and less remote from the actual data, we preferred using the latter in the main text, rather than renormalized distances.

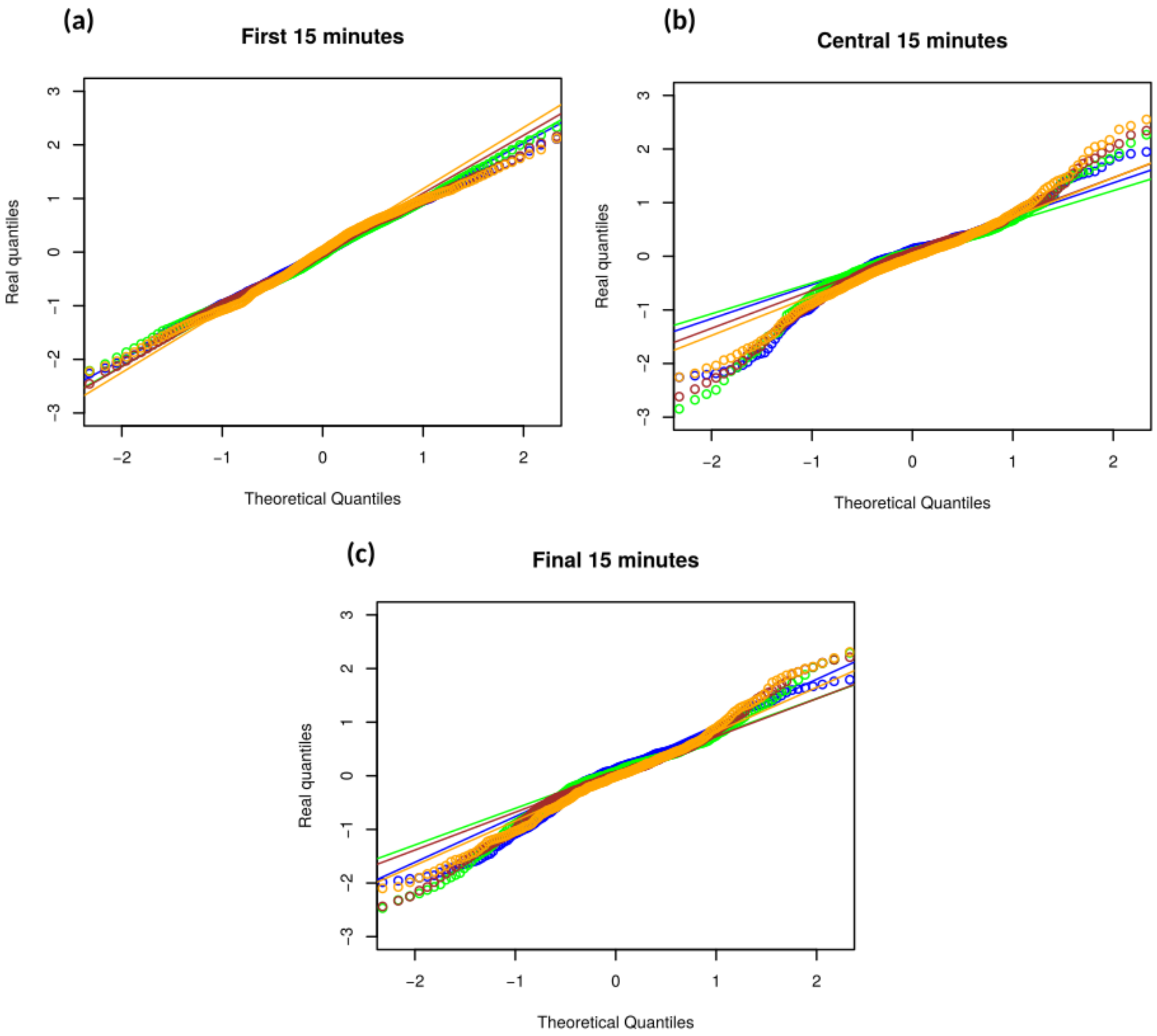


**Figure S1**: Quantile-Quantile plots of renormalized individual positions, at three different times (a: beginning, b: middle and c: end of experiments), in the four treatments (*blue*: Low Density; *green*: High Density; *brown*: Diffuse hosts; *orange*: Clumped Hosts). The solid lines are the expected relationships under Gaussian distributions. The dots are actual (observed) quantiles. Note that in all cases but the beginning of experiments (a), renormalized distributions strongly deviate from Gaussian, and are fat-tailed (remote quantiles are located further than expected under a Gaussian distribution).

#### 3. More on mixture-models fitting

The following distributions were fitted to the distribution of individuals. All distributions had mean zero (i.e. were centred on the release point) and were symmetric (even), but various free parameters controlled their shape. The first and simplest was a Normal distribution with zero mean (one free parameter: the standard deviation σ):

$$g_1(x, \sigma) = \frac{1}{\sigma\sqrt{2\pi}} exp(-\frac{1}{2}\left(\frac{x}{\sigma}\right)^2)$$

More complex models were mixtures of several Gaussian distributions, each characterised by its relative frequency and its variance. We considered a 2-mixture (3 free parameters):

$$g_2(x, \sigma_1, \sigma_2, p_1) = p_1 g_1(x, \sigma_1) + (1 - p_1) g_1(x, \sigma_2)$$

and a 3-mixture (6 free parameters):

$$g_3(x, \sigma_1, \sigma_2, \sigma_3, p_1, p_2) = p_1 g_1(x, \sigma_1) + p_2 g_1(x, \sigma_2) + (1 - p_1 - p_2) g_1(x, \sigma_3)$$

More complex Gaussian mixtures were also explored, but were not supported and/or difficult to fit to the data.
Finally, we considered the t-distribution with a rescaling factor (2 free parameters):

$$g_t(x, \sigma, \nu) = T(\frac{x}{\sigma}, \nu)$$

where *T* is the usual pdf of Student's t distribution with ν degrees of freedom.

These were in practice computed using the builtin R functions *dnorm* and *dt*. Probability densities were computed per observation, and then multiplied to produce a likelihood. Each model was thus adjusted by maximum likelihood, using function *optim* and simulated annealing as the optimization algorithm. Initial guesses for parameters were chosen as small perturbations from the simple Gaussian distribution. See the provided R code for all details. Finally, models were compared and ranked using the Akaike Information Criterion.

Overwhelmingly, both from visual inspection and AIC selection, the best fitting model was the 2-Gaussian mixture, as reported in the main text.

For the 2-Gaussian mixture model, parameter uncertainty was further quantified using bootstrap. The entire model fitting procedure was repeated B=600 times, resampling replicates. From these resamplings, the standard deviation of the distribution of bootstrap parameter estimates was used to compute a Standard Error (SE). 95% confidence intervals were then obtained for each parameter using the standard confidence interval procedure. This procedure was in this case preferred to the percentile confidence approach, as it proved more stable and required fewer resamplings, and thus less computation time, to converge.

Using the model, we would further compute the predicted proportion of resident and explorer individuals at any given place and time. This is what is used in Figure 6b in the main text. The computation is as follows. Knowing the estimated values of $(\sigma_1, \sigma_2, p_1)$, we can compute the predicted proportion of explorer individuals at some position $x$ as:

$$(1 - p_1) g_1(x, \sigma_2) / g_2(x, \sigma_1, \sigma_2, p_1)$$

##### 4. MSD of the resident component

The MSD of the explorer component, and the proportion of resident individuals are shown in Figure 4 in the main text. The MSD of the resident component is shown below in Figure S3. Its low value, and absence of clear diffusion over time, made estimations more noisy for this parameter than for the other two.

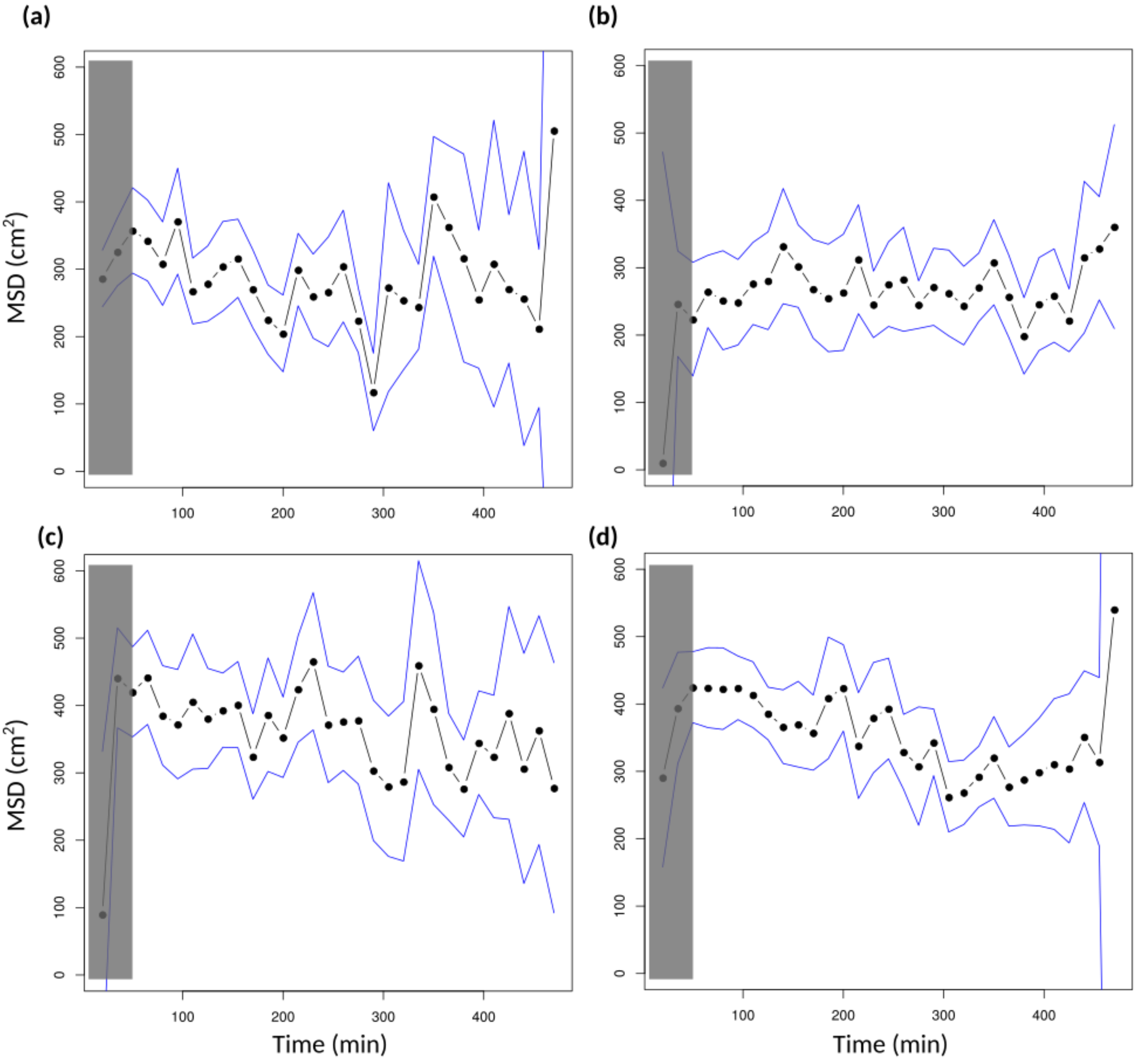


**Figure S3**: MSD of the resident components, estimated from the 2-Gaussian mixture fits, as a function of time. Solid blue lines represent 95% bootstrap confidence intervals. No regression of MSD with respect to time was significantly different from zero. As in the main Figure, the latency phase (initial 40 minutes) was shadowed, as the two components are difficult to discriminate during this phase. Note the y-scale is in $cm^2$ rather than $m^2$.

#### 5. Predictive power of different population quantiles

The correlation, at replicate level, between the advancement of the population (population quantiles) and the parasitism metrics (total parasitism rate and dispersal coefficient), was computed for a range of population quantiles, between 50% to 100%. Spearman's rank correlations were used, but similar results were obtained with Pearson's linear correlations. Results are shown below in Figure S4.

As is visible and stated in the main text, the best predictive power is achieved using around the 98% population quantile. It is thus the value used in the main text. Interestingly, 98% is also a quantile commonly recommended in population studies for the location of the population front.

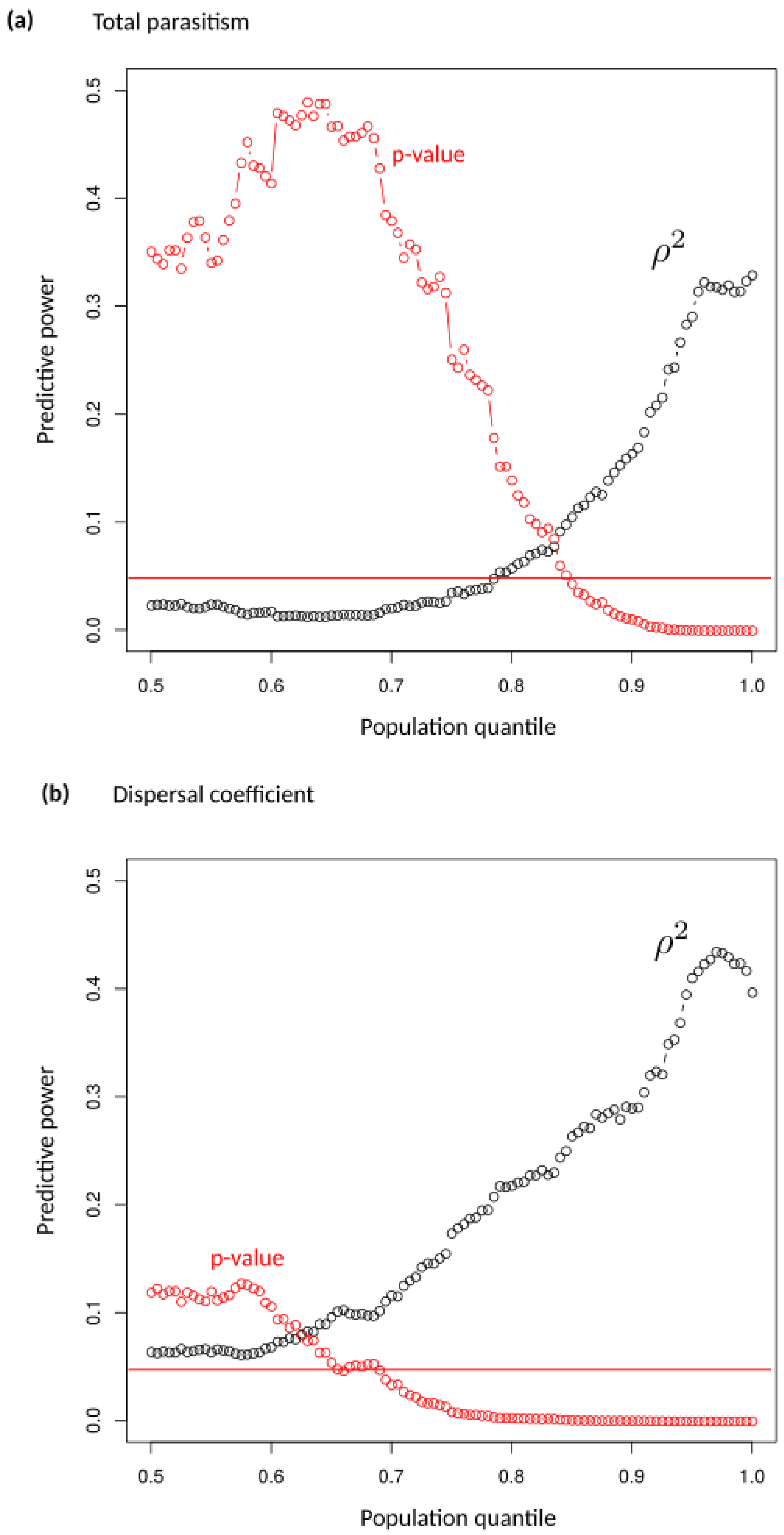


**Figure S4**: Predictive power ($\rho^2$ and p-value) of different population quantiles, with respect to total parasitism (a) and dispersal coefficient (b). Population quantiles are taken at the end of the experiments.

#### 6. Population distribution integrated over time

At a given place, i.e. for a given host, the total density of individuals that might have parasitized it is the integral over time of the population distribution. More empirically, it is simply the overall distribution of all individual detections, pooling all times together. Since the variance of the population distribution increased through time, the integrated distribution is a mixture of many distributions that have different variances. By the same argument used elsewhere in the main text, this heterogeneity of variance should create an additional leptokurticity in the pooled distribution. The pooled distribution is shown below in Fig. S5. As can be seen on the Figure, the distribution is not Gaussian either, and retains the two modes identified at any time.

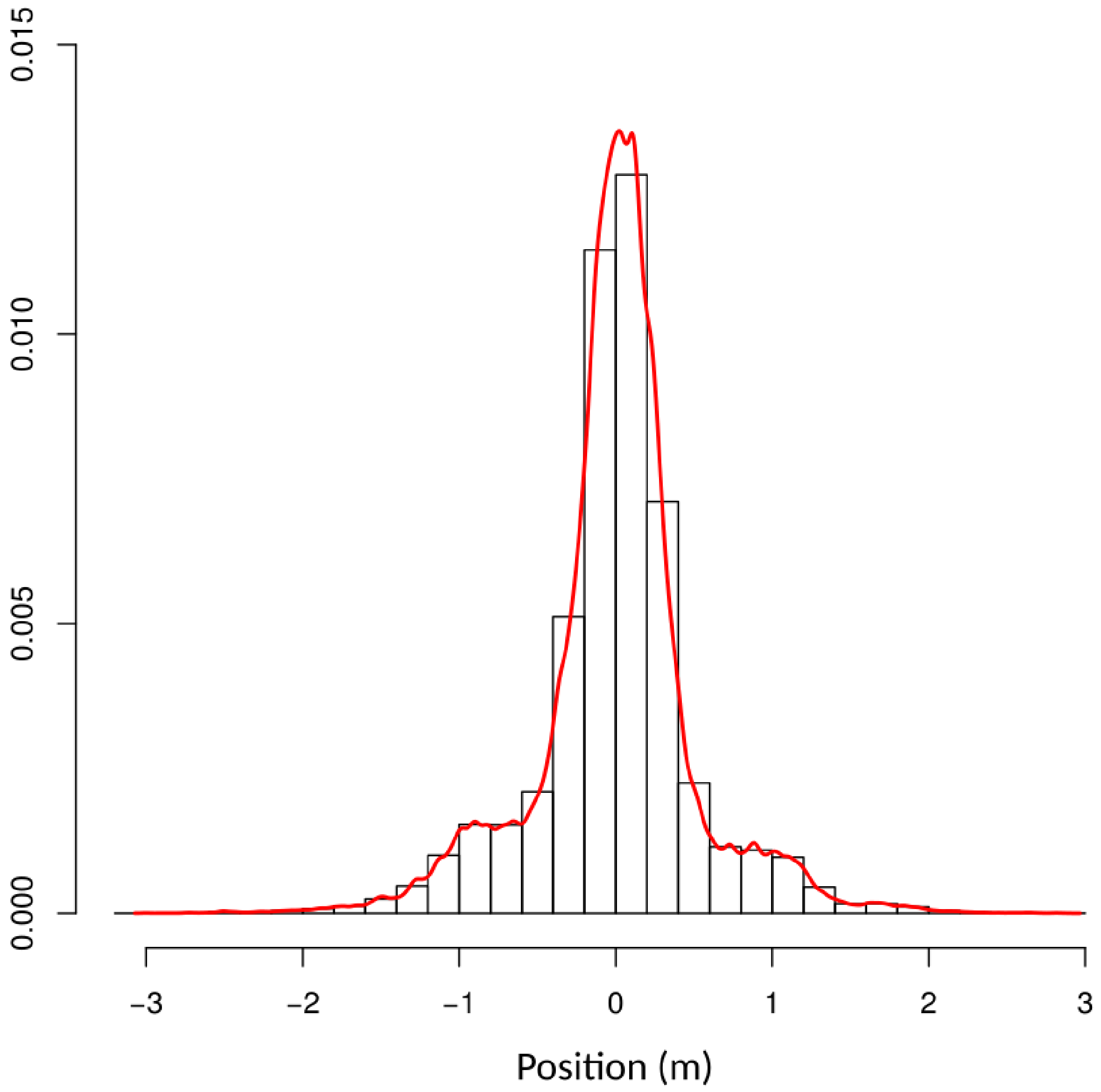


**Figure S5:** Cumulated distribution of individuals. All individual detections were pooled irrespective of time, so that spatial distributions at each time have approximately equal weight (with some variability due to the rate of detection). The red curve is a smoothed density kernel. Replicates from Diffuse and Clumped host treatments were combined in this figure (n=42 replicates).
